# Functional Exploration of Conserved Sequences in the Distal Face of Angiotensinogen

**DOI:** 10.1101/2022.12.22.521618

**Authors:** Naofumi Amioka, Chia-Hua Wu, Hisashi Sawada, Sohei Ito, Alex C. Pettey, Congqing Wu, Jessica J. Moorleghen, Deborah A. Howatt, Gregory A. Graf, Craig W. Vander Kooi, Alan Daugherty, Hong S. Lu

**Author notes:** These authors contributed equally to this work (Naofumi Amioka, Chia-Hua Wu). Corresponding Authors: Hisashi Sawada, Hong S. Lu.

## Abstract

**Background:** Angiotensinogen (AGT) is an essential component in the renin-angiotensin system. AGT has highly conserved sequences in the loop and β-sheet regions among species; however, their functions have not been studied.

**Methods:** Adeno-associated viral vector (AAV) serotype 2/8 encoding mouse AGT with mutations of conserved sequences in the loop (AAV.loop-Mut), β-sheet (AAV.βsheet-Mut), or both regions (AAV.loop/βsheet-Mut) were injected into male hepatocyte-specific AGT deficient (hepAGT-/-) mice in an LDL receptor –/– background. AAV containing mouse wild-type AGT (AAV.mAGT) or a null vector (AAV.null) were used as controls. Two weeks after AAV administration, all mice were fed a Western diet for 12 weeks. To determine how AGT secretion is regulated in hepatocytes, AAVs containing the above mutations were transducted into HepG2 cells.

**Results:** In hepAGT-/– mice infected with AAV.loop-Mut or βsheet-Mut, plasma AGT concentrations, systolic blood pressure, and atherosclerosis were comparable to those in AAV.mAGT-infected mice. Surprisingly, plasma AGT concentrations, systolic blood pressure, and atherosclerotic lesion size in hepAGT-/– mice infected with AAV.loop/βsheet-Mut were not different from mice infected with AAV.null. In contrast, hepatic *Agt* mRNA abundance was elevated to a comparable magnitude as AAV.mAGT-infected mice. Immunostaining showed that AGT protein was accumulated trol and AAV containing wild-type mouse AGT as a positive control. We have demonstrated consistently in this and previous studies tht located in the endoplasmic reticulum.

**Conclusions:** The conserved sequences in either the loop or β-sheet region individually have no effect on AGT regulation, but the conserved sequences in both regions synergistically contribute to the secretion of AGT from hepatocytes.

**HIGHLIGHTS:**

1. The loop and β-sheet regions in the distal face of angiotensinogen (AGT) have highly conserved sequences across species.
2. Mutations on either the loop or β-sheet regions do not affect plasma AGT concentrations, blood pressure, and atherosclerosis in hypercholesterolemic mice.
3. The conserved sequences in the loop and β-sheet regions regulate the secretion of AGT from hepatocytes synergistically in vivo and in cultured cells.

## INTRODUCTION

The renin-angiotensin system (RAS) exerts crucial roles in the pathophysiology of hypertension and atherosclerosis.^1–6^ Angiotensinogen (AGT), the unique substrate of the RAS, is composed of 452 amino acids in humans (453 amino acids in mice) after the cleavage of the signal peptides.^7–10^ AGT is cleaved by renin to produce angiotensin I (AngI), a decapeptide that is then cleaved to produce angiotensin II (AngII, an octapeptide) and other angiotensin peptides.^9, 10^ Although the distal portion (the non-AngI part termed as “des(AngI)AGT”) contains 98% of the amino acids of AGT, there is scant evidence regarding the role of the des(AngI)AGT domain in regulating AngII production, AngII-mediated functions, and AngII-independent effects. des(AngI)AGT has been implicated in angiogenesis.^11, 12^ Our recent study supports that des(AngI)AGT contributes to diet-induced body weight gain and liver steatosis that are independent of AngII.^13, 14^

In addition to the conserved AngI sequence among species, three-dimensional structure analyses identified highly conserved sequences in the loop (W292, S298, V299) and β-sheet (K253, H274, E422) regions of AGT from zebrafish to humans.^10, 13^ These two regions are in the des(AngI)AGT portion and are far from the renin cleavage site.^13, 15^ A computational analysis showed that these conserved sequences are exposed to the surface of AGT. Considering their high conservation and their locations in the AGT protein, we hypothesize that these conserved sequences have an impact on the functions of AGT.

In this study, we investigated the impact of the highly conserved sequences in the loop and β-sheet regions of AGT on the dynamics of AGT and AngII-mediated vascular functions. Since hepatocytes are the major source of AGT,^13, 16-19^ we used hepatocyte-specific AGT deficient (hepAGT-/-) mice. We injected adeno-associated viral vector serotype 2/8 (AAV) encoding mouse AGT with mutations of conserved sequences in the loop, β-sheet, or both regions to hepAGT-/– mice. The presence of mutations in either region alone had no effects on AGT regulation, blood pressure, and atherosclerosis in hypercholesterolemic mice. However, mutations of the conserved sequences in both regions impacted the secretion of AGT from hepatocytes.

## MATERIALS AND METHODS

Detailed materials and methods are available in the Supplemental Material. Numerical data are available in Supplemental Excel File.

### Mice

hepAGT-/– mice and their wild-type (hepAGT+/+) littermates have been developed by breeding female *Agt* floxed mice with male albumin-Cre+/0 mice as described in our previous publications.^13, 16, 17, 20, 21^ hepAGT+/+ and hepAGT-/– littermates were generated initially in a mixed 129/C57BL/6N background and were then backcrossed to C57BL/6J strain at least 6 times.^13, 16, 17, 20, 21^ All study mice were in an LDLR-/– background. Littermates were used for each experiment. Two weeks after adeno-associated viral (AAV) injections, mice were fed Western diet (Diet # TD.88137; Envigo) for 12 weeks. This study used only male mice because of the modest response of AAV encoding wild-type AGT on plasma AGT concentrations in female mice.^20^ All animal experiments in this study were performed according to protocols approved by the University of Kentucky Institutional Animal Care and Use Committee (IACUC protocol number 2006-0009 or 2018-2968).

### Production and injection of AAVs

AAVs (serotype 2/8) were produced by the Vector Core in the Gene Therapy Program at the University of Pennsylvania. Mouse *Agt* was transduced using AAV that infects predominantly hepatocytes and contained a thyroxine-binding globulin (TBG) promoter for hepatocyte-specific synthesis.^22^ AAVs containing wild-type AGT (AAV.mAGT) or an empty vector (AAV.null) were used as controls. AAV (3 × 10^10^ genome copies/mouse) were diluted in 200 μl of sterile phosphate-buffered saline (PBS) and injected into mice intraperitoneally at 6-22 weeks of age.

### Systolic blood pressure measurements

Systolic blood pressure was measured at week 11 after the injection of AAV using a non-invasive tail-cuff system (Coda 8, Kent Scientific) following our standard protocol described previously.^23^ Conscious mice were restrained in a holder and placed on a heated platform. Blood pressure was measured 20 times for three consecutive days. Data showing <60 or >250 mmHg, standard deviation >30 mmHg, or successful measurements <5 cycles were excluded.

### Quantification of atherosclerosis

Mice were euthanized using a ketamine/xylazine cocktail (90 and 10 mg/kg, respectively) at week 14 after the infection of AAV. An *en face* method was used to measure atherosclerotic lesions on the intimal surface of the thoracic aorta, as described previously.^24, 25^ The aorta was dissected and fixed in 10% neutrally buffered formalin overnight. Subsequently, perivascular tissues were removed and the intimal surface was exposed by a longitudinal cut. Aortas were then pinned on a black wax surface and *en face* images were captured using a digital camera (DS-Ri1, Nikon). All image processing and analyses were performed using a Nikon NIS-Elements software (NIS-Elements AR 5.11.0.) Atherosclerotic lesions were traced manually on the *en face* images from the ascending aorta to 3 mm distal from the orifice of the left subclavian artery. Measured data were verified independently by an investigator who was blinded to the study groups. Atherosclerotic lesion size was presented as percent lesion area as below:

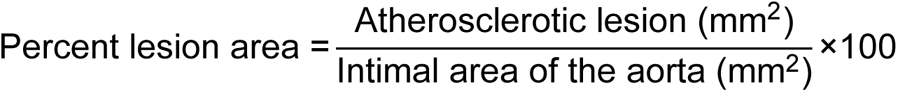

### Statistical analysis

Graphs and statistical analyses were conducted using SigmaPlot version 14.5 or 15 (SYSTAT Software Inc.) or R (version 4.1.0 lme4 package version 1.1.30). The presented data consist of the median and 25th/75th percentiles except for body weights, which were represented as mean ± SEM. Most data (except for body weight gain) were analyzed using Kruskal-Wallis one-way ANOVA on ranks, followed by Dunn’s method. The difference in body weight gain among study groups was evaluated using a linear mixed-effects model with individual mice as a random effect to account for repeated measures. Treatment group and time were included as fixed effects. Differences in slope between treatment groups were compared post-hoc with a Bonferroni correction. Statistically significant results were determined at P<0.05.

## RESULTS

### AAV-driven expression of wild-type AGT increased plasma AGT concentrations and vascular functions in hepAGT-/– mice

We first confirmed the effects of AAV encoding wild-type mouse AGT on plasma AGT concentrations, systolic blood pressure and atherosclerosis in hepAGT-/– mice. Infection of AAV.mAGT increased plasma AGT concentrations in hepAGT-/– mice that were comparable to the plasma AGT concentrations in hepAGT+/+ littermates infected with AAV containing an empty vector (**Supplemental Figure 1A**). Systolic blood pressure and atherosclerotic lesion area were increased by AAV.mAGT infection in hepAGT-/– mice to the same magnitude as in their hepAGT+/+ littermates infected with AAV.null. Plasma total cholesterol concentrations were not different among hepAGT+/+ mice infected with AAV.null, hepAGT-/– mice infected with AAV.null, and hepAGT-/– mice infected with AAV.mAGT (**Supplemental Figure 1B-D**). Therefore, in the subsequent studies, we used AAV.mAGT infection in hepAGT-/– mice to represent its wild-type control (hepAGT+/+).

### Mutations of conserved sequences in either the loop or β-sheet region of AGT increased plasma AGT concentrations and vascular functions in hepAGT-/– mice

Computational analyses revealed highly conserved sequences in the two distal face regions, the loop and β-sheet regions, of AGT (**Figure 1A**). We first determined roles of the conserved sequences in the loop region on plasma AGT concentrations, systolic blood pressure regulation and atherosclerosis. Genetic manipulation was conducted by replacing the three conserved residues in the loop region (W292, S298, V299) with alanine (**Figure 1B**). AAV encoding AGT with these mutations (AAV.loop-Mut) was injected into hepAGT-/– mice. Two weeks after the AAV injections, AAV.loop-Mut increased plasma AGT concentrations significantly in hepAGT-/– mice, which were comparable to those in hepAGT-/– mice infected with AAV.mAGT (**Figure 1C**). Plasma renin concentrations (PRC) were suppressed equivalently by the population of loop-Mut or mouse wild-type AGT (**Supplemental Figure 2A**). In comparisons with hepAGT-/– mice infected with AAV.null, infection of AAV.loop-Mut increased systolic blood pressure and atherosclerotic lesions. These phenotypes were not different from hepAGT-/– mice infected with AAV.mAGT (**Figure 1D, 1E**). Plasma total cholesterol concentrations were modestly higher in AAV.loop-Mut and AAV.mAGT-AAV infected mice, compared to AAV.null-infected mice (**Supplemental Figure 2B**).

**Figure 1.**
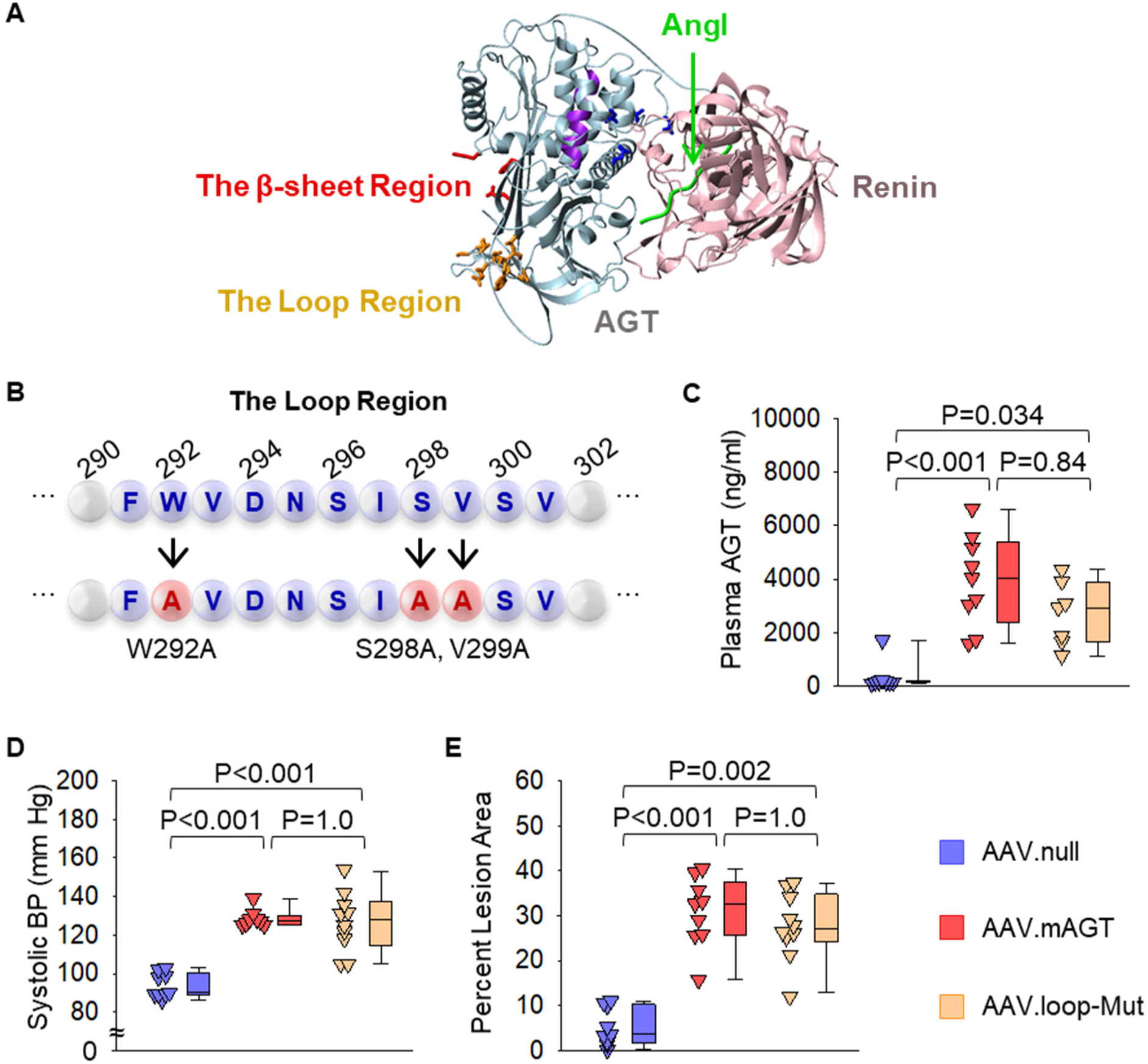
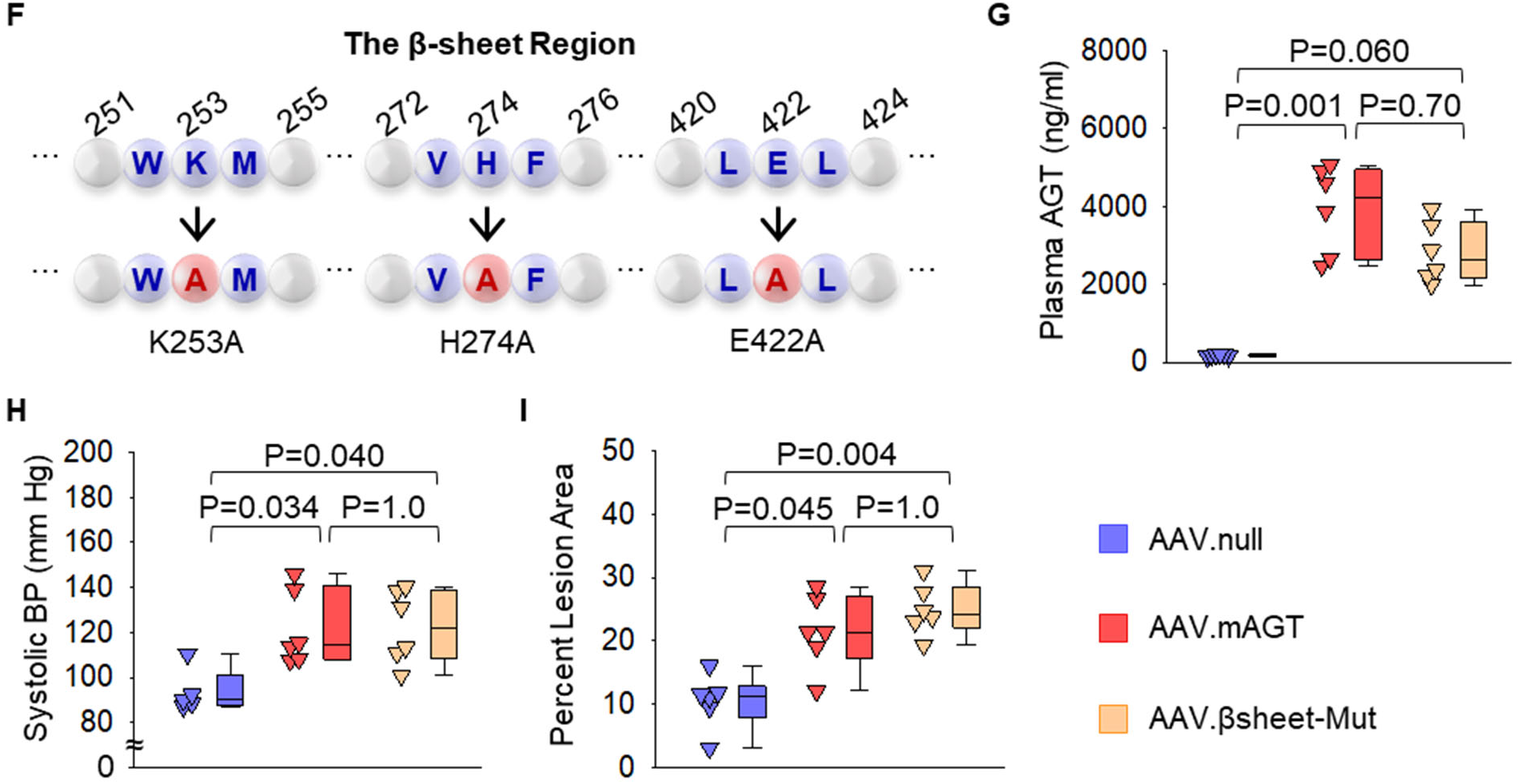
Mutations of conserved residues in either the loop or β-sheet region of AGT had comparable effects as wild-type AGT on plasma AGT concentrations, blood pressure, and atherosclerosis in hepAGT-/– mice. (**A**) Computational analysis for the three-dimensional structure of AGT and renin. The distal surface is highlighted in gold for the loop region and red for the β-sheet region. Green represents angiotensin I (AngI). (**B**) The 3 amino acid residues identical across species were replaced with alanine in the loop (W292A, S298A, V299A). (**C-E**) Male hepAGT-/– mice infected with AAVs were fed Western diet for 12 weeks. (**C**) Plasma AGT concentrations, (**D**) systolic blood pressures (BP), and (**E**) percent lesion area of atherosclerosis in hepAGT-/– mice (n = 7-10/group) infected with AAV.null, AAV.mAGT, or AAV encoding AGT with mutations in the loop region (AAV.loop-Mut). P values were determined by Kruskal-Wallis one-way ANOVA on Ranks followed by Dunn’s test. (**F**) The 3 amino acid residues identical across species were replaced with alanine in the β-sheet (K253A, H274A, E422A) regions. (**G-I**) Male hepAGT-/– mice infected with AAVs were fed Western diet for 12 weeks. (**G**) Plasma AGT concentrations, (**H**) systolic blood pressures (BP), and (**I**) percent lesion area of atherosclerosis in hepAGT-/– mice (n = 5-6/group) infected with AAV.null, AAV.mAGT, or AAV encoding AGT with mutations in the β-sheet region (AAV.βsheet-Mut). P values were determined by Kruskal-Wallis one-way ANOVA on Ranks followed by Dunn’s test.

We next determined functions of the highly conserved sequences in the β-sheet region on plasma AGT concentrations and vascular functions. The 3 highly conserved amino acid residues (K253, H274, E422) of the β-sheet region were replaced by alanine (**Figure 1F**), and AAV with mutations of these 3 residues (AAV.βsheet-Mut) were injected into hepAGT-/– mice. Injection of AAV.βsheet-Mut increased plasma AGT concentrations similar to mice injected with AAV.mAGT (**Figure 1G**). Population of βsheet-Mut also increased systolic blood pressure and atherosclerosis lesions in hepAGT-/– mice (**Figure 1H, I**). PRC was suppressed equivalently by AAV.βsheet-Mut and AAV.mAGT (**Supplemental Figure 3A**). Plasma total cholesterol concentrations were not different among the 3 groups (**Supplemental Figure 3B**).

We have observed consistently that hepAGT-/– mice have reduced body weight gain and attenuated liver steatosis when fed Western diet.^13, 14^ Expression of wild-type AGT mutations of conserved sequences in either the loop or β-sheet region of AGT to hepAGT-/– mice led to increased body weight gain and liver weight, compared to those in hepAGT-/– mice infected with AAV.null (**Supplemental Figures 4 and 5**).

### Mutations of the conserved residues in both the loop and β-sheet regions did not change plasma AGT concentrations and vascular functions in hepAGT-/– mice

We next generated AAV encoding mouse AGT with mutations on the conserved amino acid residues of both the loop and β-sheet regions (AAV.loop/βsheet-Mut). The computational analysis did not detect major structural changes by the mutations. Unlike AGT with mutations in either region, plasma AGT concentrations were not increased (**Figure 2A**) and PRC was not altered (**Supplemental Figure 6A**) by AAV.loop/βsheet– Mut infection in hepAGT-/– mice. Plasma total cholesterol concentrations did not differ among the 3 groups (**Supplemental Figure 6B**). AAV.loop/βsheet-Mut did not increase systolic blood pressure or augment atherosclerotic lesion size (**Figure 2B and C**). These results suggest that the conserved sequences in the loop and β-sheet regions have synergistic effects on plasma AGT concentrations and vascular functions.

**Figure 2.**
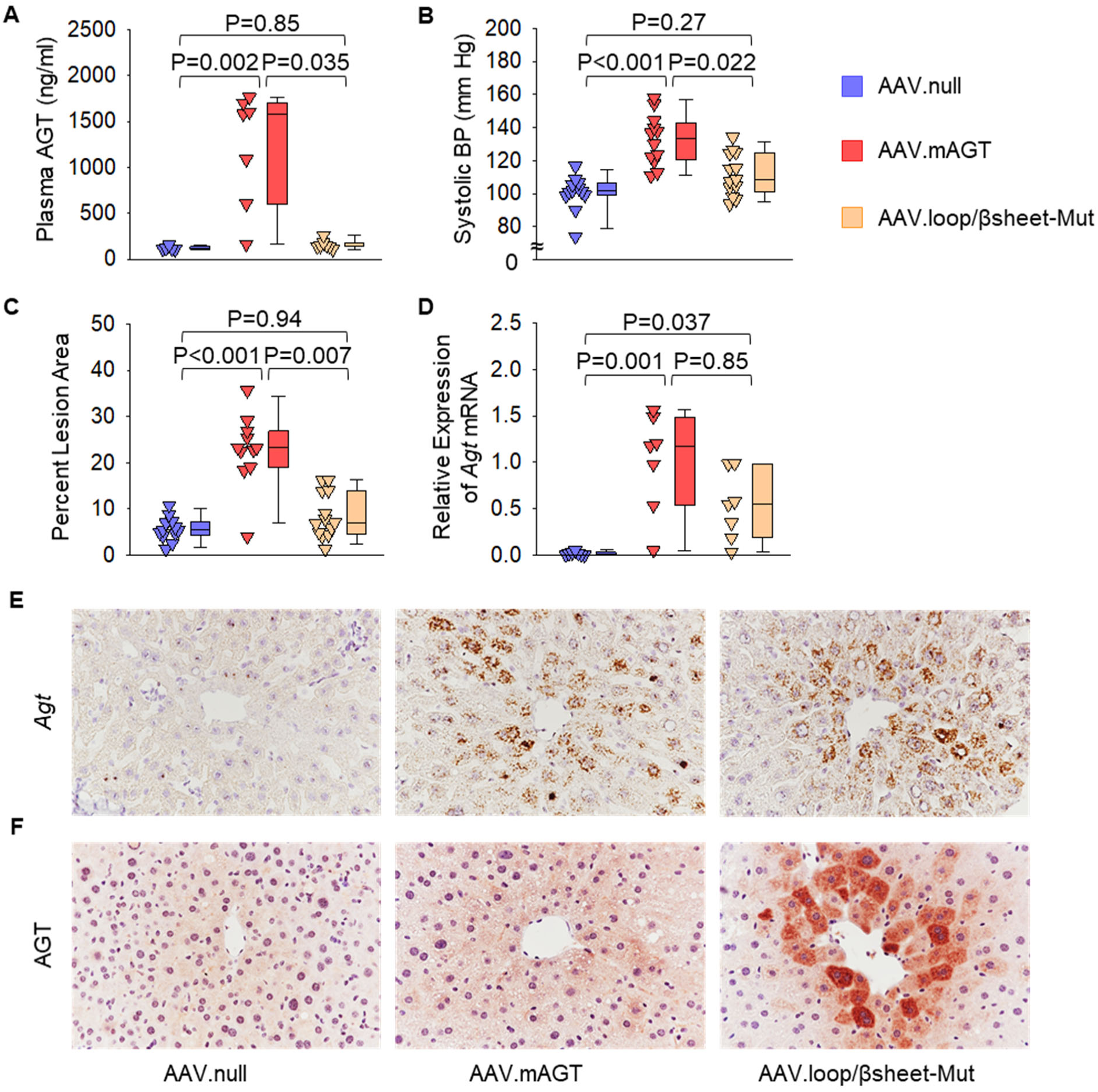
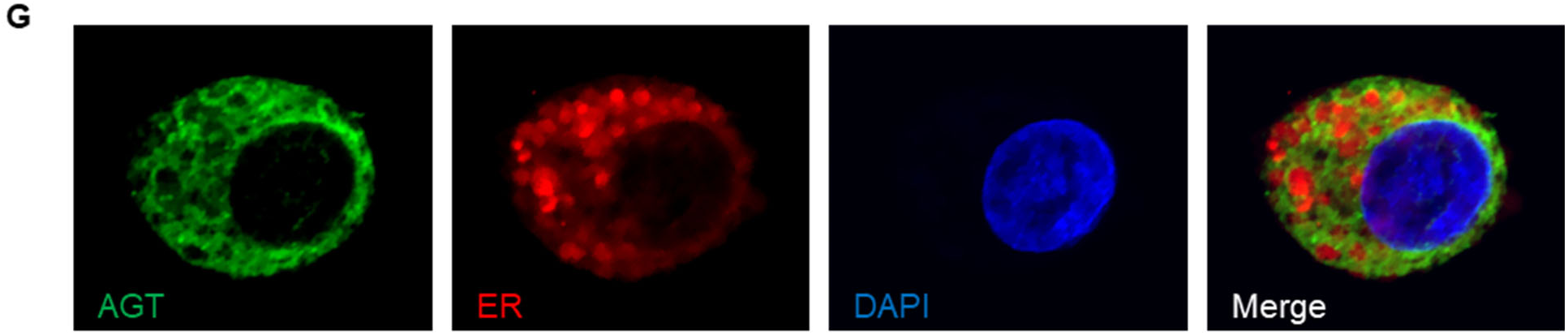
Mutations of conserved residues in both the loop and β-sheet regions of AGT led to AGT accumulation in hepatocytes. (**A-F**) Male hepAGT-/– mice infected with AAV were fed Western diet for 12 weeks. (**A**) Plasma AGT concentrations, (**B**) systolic blood pressures (BP), (**C**) percent lesion area of atherosclerosis, (**D**) *Agt* mRNA abundance in liver, (**E**) RNAscope of *Agt* mRNA in liver, and (**F**) immunostaining of AGT in liver of hepAGT-/– mice infected with AAV.null, AAV.mAGT, or AAV.loop/βsheet-Mut (n = 5-13/group). P values were determined by Kruskal-Wallis one-way ANOVA on Ranks followed by Dunn’s test. (**G**) Co-staining of AGT and ER (endoplasmic reticulum) in cultured HepG2 cells transducted with AAV.loop/βsheet-Mut.

Liver is the primary source for AGT production,^13, 16-18^ and the AAVs we used predominantly populate AGT in hepatocytes.^22^ Therefore, we examined the abundance of hepatic *Agt* mRNA by qPCR. Hepatic *Agt* mRNA was comparable between mice infected with AAV.loop/βsheet-Mut and AAV.mAGT (**Figure 2D**). A previous study reported that AAV predominantly leads to infection in pericentral hepatocytes.^22^ Consistently, in situ hybridization by RNAscope revealed the presence of *Agt* mRNA in hepatocytes, predominantly in pericentral hepatocytes of both AAV.loop/βsheet-Mut– and AAV.mAGT-infected mice, but *Agt* mRNA was not detected in the liver of AAV.null-infected mice (**Figure 2E**). We have not been able to detect AGT protein in hepatocytes in hepAGT+/+ mice under normal conditions using immunostaining, although it has been found consistently in renal proximal tubule cells (**Supplemental Figure 7**). As expected, AGT protein was not detected in hepatocytes of AAV.mAGT-infected mice (**Figure 2F**). However, AGT protein was accumulated in hepatocytes of mice infected with AAV.loop/βsheet-Mut, as detected by immunostaining (**Figure 2F**). Accumulation of AGT protein was predominantly in pericentral hepatocytes as confirmed with immunostaining of glutamine synthetase, an enzyme that is abundant in hepatocytes surrounding central veins (**Supplemental Figure 8**).

To determine whether des(AngI)AGT, the remaining part of AGT after cleavage of AngI by renin, impairs its secretion from liver, we measured plasma AGT concentrations in hepAGT-/– mice infected with AAVs containing des(AngI)AGT (AAV.des(AngI)AGT) alone. As shown in **Supplemental Figure 9**, population of des(angI)AGT led to plasma AGT concentrations more than 20-fold higher in hepAGT-/– mice.

Despite the profound accumulation of AGT intracellularly, expression of AAV.loop/βsheet-Mut did not increase body weight gain or liver weight, compared to AAV.null in hepAGT-/– mice (**Supplemental Figure 10A and B**). Liver steatosis, as shown by histology of liver sections, was evident in hepAGT-/– mice infected with wild type AGT (AAV.mAGT), but not those infected with AAV.loop/βsheet-Mut (**Supplemental Figure 11**).

### AGT with mutations in both the loop and β-sheet regions did not accumulate in endoplasmic reticulum of HepG2 cells

To determine the loci of accumulation of AGT with mutations in both the loop and β– sheet regions, HepG2 cells were transducted with AAV.null, AAV.mAGT, AAV.loop-Mut, AAV.βsheet-Mut, or AAV.loop/βsheet-Mut. Consistent with in vivo studies, mAGT was not detected in HepG2 cells transducted with either wild-type AGT or mutations in either the loop or β-sheet region. In contrast, transduction of AAV.loop/βsheet-Mut led to AGT accumulation in HepG2 cells. To determine the cellular location of accumulation of mutated AGT, we simultaneously stained AGT and ER (endoplasmic reticulum) that is important for protein processing and secretion.^26–28^ Confocal microscopy revealed that AGT immunostaining did not overlapped with ER staining (**Figure 2G**).

### AGT with mutations in both the loop and β-sheet regions did not impair glycosylation of AGT in liver

Post-translational modifications, such as glycosylation, play a pivotal role in protein maturation and secretion.^29^ AGT has multiple potential glycosylation sites.^9, 30^ Therefore, we determined whether mutated AGT impaired glycosylation. As shown by Western blotting, AGT with the β-sheet mutations had comparable molecular weight of wild-type AGT, while AGT with loop mutants alone or mutants in both regions exhibited higher molecular weights (∼2 kDa higher). After deglycosylation, Western blotting showed that the molecular weights of wild-type and any of the AGT mutations were comparable (**Supplemental Figure 12**).

## DISCUSSION

Our study is the first to determine whether the highly conserved sequences in the distal regions of AGT are functionally important for AGT regulation. We used mice in which the *Agt* gene was deleted specifically in hepatocytes. Mouse AGT containing mutations of the conserved residues in the loop, β-sheet, or both regions were populated to the liver of this mouse model by an AAV vector. This AAV serotype 2/8 vector containing the TBG promoter targets hepatocytes with high specificity.^22, 31^ We used AAV containing a null vector as a negative control and AAV containing wild-type mouse AGT as a positive control. We have demonstrated consistently in this and previous studies that the genome copies of AAV encoding mouse wild-type AGT in hepAGT-/– mice led to comparable plasma AGT concentrations and AngII-mediated vascular functions such as blood pressure and atherosclerosis with those in hepAGT+/+ mice.^13, 16, 20^ Comparable to hepAGT-/– mice infected with AAV.mAGT, plasma AGT concentrations were increased in hepAGT-/– mice infected with AAV.loop-Mut or AAV.βsheet-Mut alone, but not in hepAGT-/– mice infected with AAV.loop/βsheet-Mut. It is clear that mutations of conserved sequences in either region do not affect AGT regulation and AGT-mediated vascular functions, but the conserved sequences of both regions in combination are needed to maintain the normal regulation of AGT. Overall, these data suggest that the des(AngI)AGT part of AGT contributes to AGT regulation.

mRNA abundance of AGT in liver of hepAGT-/– mice infected with AAV.loop/βsheet-Mut was equivalent to its abundance in liver of hepAGT-/– mice infected with AAV.mAGT, supporting that the mutations of the conserved sequences do not affect AGT mRNA expression. Under normal conditions, AGT is produced and secreted rapidly from hepatocytes to plasma. Therefore, it is technically difficult to detect AGT protein in hepatocytes by immunostaining. In contrast, we found that high abundance of AGT protein, as demonstrated by immunostaining, was accumulated in hepatocytes surrounding central veins in liver sections of hepAGT-/– mice infected with AAV.loop/βsheet-Mut. In agreement with the in vivo data, AGT protein accumulated in cultured HepG2 cells transducted with AAV.loop/βsheet-Mut. These findings implicate that the conserved sequences in the β-sheet and loop regions of AGT do not affect mRNA or protein production, but impair the secretion of AGT protein from hepatopcytes. Consequently, plasma AGT concentration remained low, and systolic blood pressure and atherosclerosis were not changed in hepAGT-/– mice infected with AAV.loop/βsheet-Mut.

AGT is cleaved by renin to produce AngI and des(AngI)AGT. As explored by Yan and colleagues,^32^ AGT binds renin through an allosteric mechanism that directly interacts with the N-terminus of AGT. Therefore, conserved sequences in the des(AngI)AGT domain of AGT that contains both the loop and the β-sheet regions should not be associated with AngI release from AGT. Deletion of AGT in hepatocytes attenuated Western diet-induced body weight gain and liver steatosis that is attributed to des(AngI)AGT.^13, 14^ des(AngI)AGT can be readily secreted into the circulation, as shown by high plasma AGT concentrations in hepAGT-/– mice infected with AAVs containing des(AngI)AGT. Mutations of the conserved sequences in either the loop or the β-sheet region did not affect body weight gain or liver weight, implicating that neither region alone affects the des(AngI)AGT function. Although mutations in both the loop and β– sheet regions of AGT led to AGT accumulation intracellularly, the mutations did not increase body weight gain or liver steatosis. It is possible that mutated AGT is not functional. It is also possible that these metabolic phenotypes are not attributed to an AGT intracellular mechanism. The present study is not able to determine whether the conserved sequences in the distal face of AGT are required to maintain des(AngI)AGT function on diet-induced body weight gain and liver steatosis due to the impaired secretion of the AGT protein. The definitive mechanism driving these phenotypes remain to be clarified in future studies.

The members of the Serpin superfamily including AGT share a unique structure containing an active loop and β-sheet. Mutations in α1-antitrypsin, a member of the serpin superfamily, lead to its accumulation in the ER of hepatocytes, resulting in liver dysfunction in humans.^33^ Locations of α1-antitrypsin mutations correspond to K337 and A401 in AGT. The conserved sequences of AGT locate at W292, S298, V299, K253, H274, and E422. Thus, the locations of the AGT mutations in the present study are different from those of α1-antitrypsin mutations. Of note, the intracellular accumulation of mutated AGT was not observed in the ER as demonstrated by HepG2 cell culture experiments. Therefore, AGT intracellular accumulation may be driven by a different mechanism from α1-antitrypsin mutations.

AGT was accumulated ubiquitously in HepG2 cells transducted with AAV.loop/βsheet-Mut. Unfortunately, simultaneous-staining of AGT and Golgi apparatus was not successful because permeabilization of cell membranes using detergent was needed for AGT staining, but it impaired staining of Golgi apparatus. Based on the diffused distribution of AGT in hepatocytes, it is unlikely to be in Golgi apparatus. It is possible that AGT is trapped in other organelles such as secretory vesicles. We have not been able to validate any authentic markers for secretory vesicles in hepatocytes. Western blotting of liver tissues supported the notion that mutations of either a single region or both regions did not impair glycosylation of AGT. It is likely AGT glycosylation is not a key determinant factor for accumulation of loop/βsheet-Mut AGT in hepatocytes. Although mutated AGT may not be trapped in either ER or Golgi apparatus, and it does not impact glycosylation, we were unable to distinguish the definitive mechanisms by which that the conserved sequences in the distal regions of AGT contributes to the secretion of AGT. The computational analysis did not support the notion that mutations of the conserved sequences in both regions affect the AGT structure; however, this has not been able to be confirmed using an in vivo approach.

There is compelling evidence that the RAS exerts a critical role in the pathophysiology of cardiovascular diseases including metabolic disorder-induced cardiovascular dysfunction.^34^ A large number of studies have been conducted to investigate their physiological and pathophysiological functions. AGT, as the sole substrate at the top of the RAS, is a unique target to regulate the RAS. Many studies primarily focused on investigating the regulation of the N-terminus of AGT, AngII. The present study is the first to report the function of this distal domain of AGT on its secretion from hepatocytes. Recently, pharmacological silencing of hepatocyte AGT has gained attention as a novel and highly potent therapy for hypertension and atherosclerosis.^13, 35, 36^ Therefore, providing a deeper understanding of AGT biology including the regulation of its secretion from hepatocytes may have a clinical impact on the optimization of targeting liver AGT as a favorable therapy for cardiovascular disease.

## DISCLOSURE

None.

## Non-standard Abbreviations

AAV: adeno-associated viral vector
AGT: angiotensinogen
AngI: angiotensin I
AngII: angiotensin II
RAS: renin-angiotensin system

## SUPPLEMENTAL MATERIALS

### MATERIALS AND METHODS

#### Computational analysis of protein structure

The structure of angiotensinogen^1^ bound to renin (PDB 2X0B) and angiotensinogen (PDB 2WXX) were utilized for molecular visualization with Pymol (The PyMOL Molecular Graphics System Version 2.1 Schrödinger; LLC).

#### Mice

Bedding was provided by P.J. Murphy (Coarse SaniChip) and changed weekly. Cotton pads were provided as enrichment. The room temperature and humidity were maintained at 68 to 74°F and 50 to 60%, respectively. Mice were maintained on a 14:10 hour light:dark cycle. Genotypes were determined by polymerase chain reaction (PCR) at weaning and termination. DNA was extracted from tails using Maxwell DNA purification kits (Cat # AS1120; Promega). The information on primers for PCR is shown in **Major Resources Table**.

#### Plasma angiotensinogen (AGT) concentrations

Blood samples for measuring plasma AGT concentrations were collected by retro-orbital bleeding with EDTA (0.5 M) at 2 weeks after the injection of AAVs. In hepAGT-/– mice infected with AAV.des(AngI)AGT, blood samples were collected at 14 weeks after the injection of the AAVs. Collected samples were centrifuged at 400 g × 20 minutes at 4°C to separate plasma. Plasma AGT concentrations were measured using mouse AGT ELISA kits (Cat # 27413; IBL America).

#### Plasma renin concentrations

Blood was collected at termination by cardiac bleeding via the right ventricle into EDTA (0.5 M) for measuring plasma renin concentrations. Plasma renin concentrations were measured using mouse renin ELISA kits (Cat # DY4277; R&D Systems) in samples supplemented with exogenous mouse recombinant AGT.

#### Plasma total cholesterol concentrations

Plasma total cholesterol concentrations at termination were measured using enzymatic kits (Cat # 999-02601; FUJIFILM Medical Systems USA).

#### RNA isolation and quantitative polymerase chain reaction (qPCR)

RNA was extracted from livers using Maxwell^®^ 16 LEV simplyRNA Purification kits (Cat # AS1280; Promega) in accordance with the manufacturer’s protocol. RNA was transcribed reversely to cDNA using iScript™ cDNA Synthesis kits (Cat # 170-8891; Bio-Rad). Quantitative PCR was performed to quantify *Agt* mRNA abundance in liver using SsoFast™ EvaGreen® Supermix kits (Cat # 172-5204; Bio-Rad) on a Bio-Rad CFX96 cycler. Data were analyzed using ΔΔCt method and normalized by the geometric mean of 3 reference genes: *Actb*, *Gapdh*, and *Rplp2*.

#### Immunostaining

Immunostaining was performed using Xmatrx® Infinity, an automated staining system (Cat #: AS4000RX; BioGenex), on paraffin-embedded sections to determine the distribution of AGT and glutamine synthetase in the liver. After fixation using paraformaldehyde (4% wt/vol), liver and kidney samples were incubated in ethanol (70% vo/vol) for 24 hours, embedded into paraffin, and cut at a thickness of 4 µm. Subsequently, sections were deparaffinized using limonene (Cat # 183164; Millipore-Sigma) followed by 2 washes with absolute ethanol (Cat # HC-800-1GAL; Fisher Scientific), and 1 wash with automation buffer (Cat # GTX30931; GeneTex). Deparaffinized sections were incubated with H_2_O_2_ (1% vol/vol; Cat # H325-500; Fisher Scientific) for 10 minutes at room temperature and then antigen retrieval (Cat # HK547-XAK; BioGenex) for 20 minutes at 98°C. Non-specific binding sites were blocked using goat serum (2.5 % vol/vol; Cat # MP-7451; Vector laboratories) for 20 minutes at room temperature. Sections were then incubated with rabbit anti-AGT antibody (0.3 µg/ml, Cat # 28101; IBL America) diluted in primary antibody diluent (Cat #: GTX28208; GeneTex) for 12 hours at 4°C or diluted rabbit anti-glutamine synthetase antibody (1.0 µg/ml, Cat # ab73593; abcam) for 15 min at 40°C. Goat anti-rabbit IgG conjugated with horseradish peroxidase (30 minutes, Cat # MP-7451; Vector laboratories) was used as the secondary antibody. ImmPACT® NovaRed (Cat # SK4805; Vector) was used as a chromogen, and hematoxylin (Cat # 26043-05; Electron Microscopy Sciences) was used for counterstaining. Slides were coverslipped with mounting medium (Cat # H-5000; Vector). Histological images were captured using an Eclipse Ni microscope at 40× magnification.

For immunostaining of HepG2 cells, cells were fixed with paraformaldehyde (4% wt/vol) for 10 minutes at 37°C and then permeabilized with Triton X-100 in PBS (0.1% vol/vol) for 1 minute at room temperature. To visualize ER (endoplasmic reticulum), we incubated cells with Red Detection Reagent in Cytopainter kits (Cat # ab139482; abcam) for 30 minutes at 37°C. After blocking with normal goat serum (Cat # MP-7451; Vector Laboratories) for 30 minutes at room temperature, the cells were incubated at 4°C with primary antibodies for AGT (Cat # 28101; IBL America) diluted in primary antibody diluent overnight. AGT was detected by incubating the cells with a fluorescent-conjugated goat anti-rabbit secondary antibody (Cat # ab150077; abcam), washing, and mounting them using an anti-fade reagent with DAPI (Cat # ab104139; abcam). Fluorescent images were captured at 100× magnification using a Nikon A1R inverted confocal microscope.

#### Hematoxylin and eosin staining

Paraffin-embedded liver sections were stained with hematoxylin and eosin. After removal of paraffin, sections were stained with eosin (Cat # ab246824, abcam) for 2 minutes and then rinsed with automation buffer. Subsequently, sections were stained with hematoxylin for 30 seconds, rinsed with automation buffer and water, and allowed to air dry. Finally, slides were coverslipped with mounting medium. Images were captured at 10× magnification using an Eclipse Ni microscope.

#### In situ hybridization

The distribution of *Agt* mRNA in liver tissue sections was examined by RNAscope® following the manufacturer’s instructions (Advanced Cell Diagnostics). Briefly, after deparaffinization using xylene (Cat # 89370-088; VWR), sections were incubated with H_2_O_2_ for 10 minutes at room temperature. Then, target retrieval (Cat # 26043-05; Advanced Cell Diagnostics) was performed for 30 minutes at 100°C, followed by a protease (Cat # 322331; Advanced Cell Diagnostics) incubation step for 15 minutes at 40 °C. Target mRNA was hybridized with mouse *Agt* probe (Cat # 426941; Advanced Cell Diagnostics) for 2 hours at 40°C, and amplified signals were detected using diaminobenzidine (Cat # 322310; Advanced Cell Diagnostics). Hematoxylin was used to stain nuclei.

#### Western blotting

Liver tissues harvested from study mice were homogenized in RIPA buffer (Cat #9803; Cell Signaling Technology) with protease inhibitor cocktail (Cat #P8340; MilliporeSigma). Protein concentrations were determined using DC assay kits (Cat # 5000111; Bio-Rad). Equal amounts of protein samples (20 µg) were resolved by SDS-PAGE (10% wt/vol) and transferred electrophoretically to PVDF membranes (Cat #1704156 or #1704273; Bio-Rad). After blocking, antibodies against the following proteins were used to probe membranes: AGT (Cat # 28101; IBL America) and β-actin (Cat # A5441, MilliporeSigma). Membranes were incubated with goat anti-rabbit (Cat # PI-1000, Vector Laboratories) to detect AGT and goat anti-mouse secondary antibodies (Cat # A2554, MilliporeSigma) to detect β-actin, respectively. For Western blot of deglycosylated proteins, Protein Deglycosylation Mix II (Cat # P6044; NEB) was used following the manufacturer’s protocol. In brief, after mixed with H_2_O (40 μl) and 10× Deglycosylation Mix Buffer 2 (5 μl), proteins (100 μg) were incubated at 75°C for 10 minutes and then cooled down. Subsequently, the protein mix was incubated with Protein Deglycosylation Mix II at room temperature for 30 minutes and then incubated at 37°C for 1 hour. Finally, deglycosylated protein (20 µg) was used for Western blot.

#### Cell culture

HepG2 (Passages 12-17, ATCC) were cultured in Dulbecco’s Modified Eagle Medium (DMEM) supplemented with High Glucose (Cat # 12100046; Gibco), FBS (10% vol/vol; R&D), penicillin-streptomycin (1% vol/vol; Cat # 15070063; Gibco), and GlutaMax I (1% vol/vol; Cat # 35050061; Gibco) under a humidified atmosphere of CO_2_ (5% at 37°C. Prior to AAV transduction, cells were seeded onto 4-well Lab-Tek® II Chamber Slide™ (1.0 × 10^4^ cells/well, Cat # 154526; Thermo Scientific) and allowed to grow for 24 hours.

To optimize the multiplicity of infection (MOI) of AAVs in HepG2 cells, AAVs conjugating with EGFP (enhanced green fluorescent protein) driven by the TBG (thyroxine-binding globulin) promoter was transducted into HepG2 cells with one of the three MOIs (1.0 × 10^5^, 1.0 × 10^6^, or 1.0 × 10^7^). EGFP was detected in less than 3% of cells transducted with AAVs of 1.0 × 10^5^ or 1.0 × 10^6^ MOI, while AAVs of 1.0 × 10^7^ MOI provided ∼10% transduction into HepG2 cells (as shown in the image below). This result is consistent with a previous report using the same AAV serotype and the same promoter in HepG2 cells.^2^ Therefore, in experiments using HepG2 cells, we used the MOI of 1.0 × 10^7^ for AAVs containing AGT or its mutants.

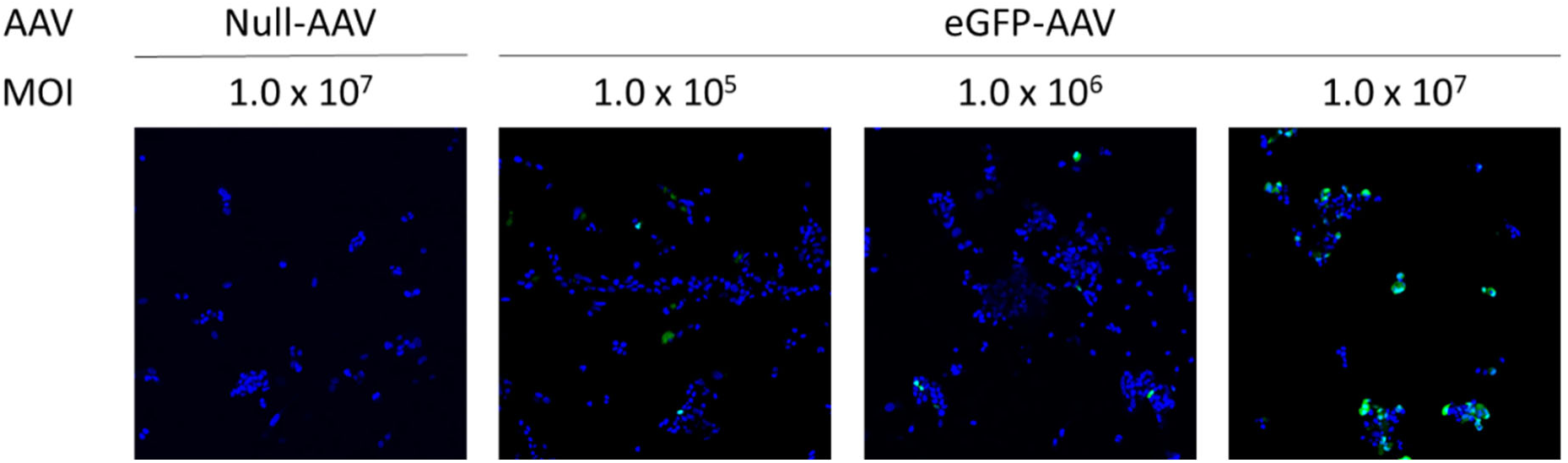

## MAJOR RESOURCES TABLES

Animals (in vivo studies) – Mice

Animals (in vivo studies) - Mice
| Genotypes | Sex | Vendor or Source | Persistent ID / URL |
| --- | --- | --- | --- |
| *Agt* f/f albumin-*Cre* 0/0 LDL receptor -/- | Male | In house breeding | N/A |
| *Agt* f/f albumin-*Cre* +/0 LDL receptor -/- | Male | In house breeding | N/A |

Breeders – Mice

Breeders – Mice
| Genotypes | Sex | Background Strain | Vendor or Source | Persistent ID / URL |
| --- | --- | --- | --- | --- |
| *Agt* f/f albumin-*Cre* +/0 LDL receptor -/- | Male | C57BL/6J (N≥6) | In house breeding | N/A |
| *Agt* f/f albumin-*Cre* 0/0 LDL receptor -/- | Female | C57BL/6J (N≥6) | In house breeding | N/A |

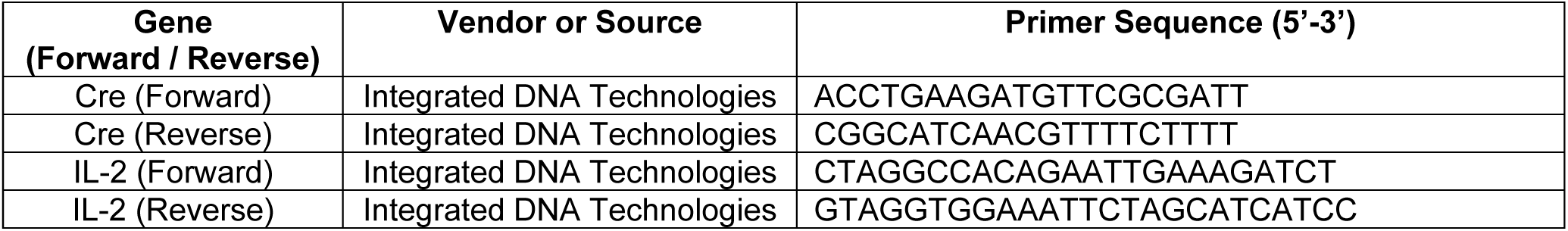
Primer sequences for genotyping of Cre

Reagents for RNAscope

Reagents for RNAscope
| Reagents | Vendor or Source | Catalog # | Lot # | Persistent ID / URL |
| --- | --- | --- | --- | --- |
| Retrieval | Advanced Cell Diagnostics | 322001 | 2014520 | N/AMAJOR RESOURCES |
| Proteinase | Advanced Cell Diagnostics | 322331 | 2011935 | https://acdbio.com/store/rnascope-protease-plus.html |
| Probe (Mm *Agt*) | Advanced Cell Diagnostics | 426941 | 20244A | N/A |
| RNAscope 2.5 HD Detection Reagents Brown | Advanced Cell Diagnostics | 322310 | N/A | N/A |

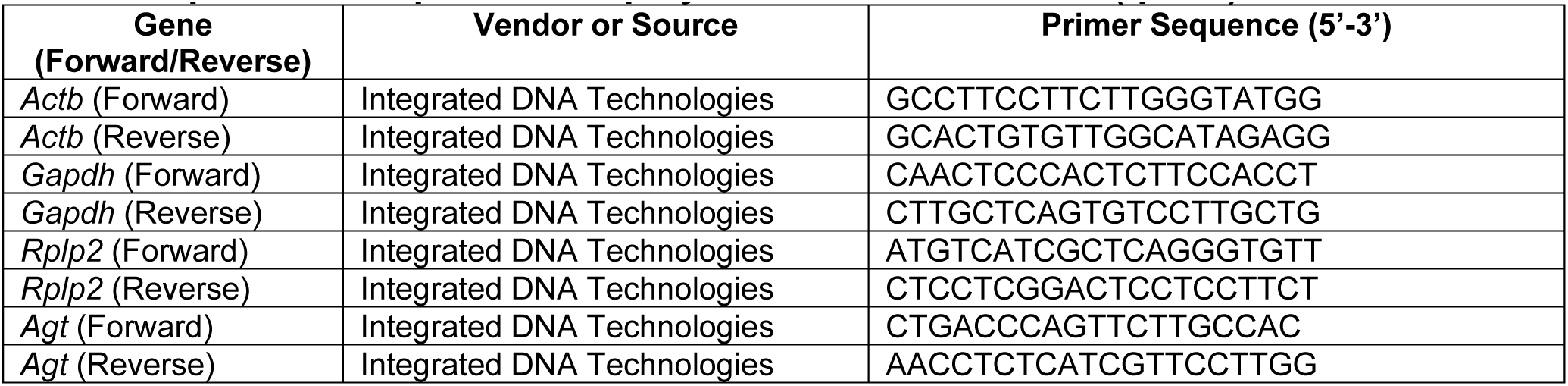
Primer sequences for quantitative polymerase chain reaction (qPCR)

Primary antibodies

Primary antibodies
| Target antigen | Vendor | Catalog # | Working concentration | Lot # | Persistent ID / URL |
| --- | --- | --- | --- | --- | --- |
| Mouse angiotensinogen | Immuno-Biological Laboratories | 28101 | 0.3 µg/ml<br>(immunostaining - liver)<br>1.0 µg/ml<br>(immunostaining – hepG2 cells)<br>1.0 µg/ml<br>(Western blot - liver) | 11-711 | https://www.ibl-america.com/angiotensinogen-405-anti-mouse-rat-rabbit-igg-affinity-purify-1/ |
| Glutamine synthetase | Abcam | ab73593 | 1.0 µg/ml<br>(immunostaining) | GR3374<br>144 | https://www.abcam.com/glutamine-synthetase-antibody-ab73593.html |
| β-actin | MilliporeSigma | A5441 | 1:3000<br>(Western blot) | 029M48<br>83V | https://www.sigmaaldrich.com/US/en/product/sigma/a5441 |

Materials for immunostaining

Materials for immunostaining
| Reagents | Vendor or Source | Catalog # | Lot # | Persistent ID / URL |
| --- | --- | --- | --- | --- |
| Immuno Automation Buffer™ | GeneTex | GTX30931 | 822203993 | https://www.genetex.com/Product/Detail/Immuno-Automation-Buffer-trade-10X/GTX30931 |
| EZ-ARTM 2 Elegance | BioGenex | HX032YCX-GP | HK5471220 | https://biogenex.com/product/ez-ar-2-elegance-edta/?v=7516fd43adaa |
| ImmPRESS® HRP Goat Anti-Rabbit IgG Polymer Detection Kit, Peroxidase | Vector Laboratories | MP-7451 | ZG0723 | https://vectorlabs.com/products/enzyme-polymer/immpress-hrp-goat-anti-rabbit-igg-kit |
| Primary Antibody Diluent | GeneTex | GTX28208 | 822203742 | https://www.genetex.com/Product/Detail/Primary-Antibody-Diluent/GTX28208 |
| ImmPACT NovaRED® Substrate Kit, Peroxidase (HRP) | Vector Laboratories | SK-4805 | ZH0128 | https://vectorlabs.com/products/substrates/impact-novared-hrp-substrate |
| Mayer's Hematoxylin | Electron Microscopy Sciences | 26043-05 | 220831-18 | https://www.emsdiasum.com/mayers-hematoxylin-500ml |
| VectaMount® Permanent Mounting Medium | Vector Laboratories | H-5000 | U0122 | https://vectorlabs.com/products/mounting/vectamount-permanent-mounting-medium |
| Cytopainter ER | abcam | ab139482 | 1009102-3 | https://www.abcam.com/products/assay-kits/er-staining-kit-red-fluorescence-cytopainter-ab139482.html |
| Goat Anti-Rabbit IgG H&L (Alexa Fluor® 488) | abcam | ab150077 | GR297743-1 | https://www.abcam.com/products/secondary-antibodies/goat-rabbit-igg-hl-alex-fluor-488-ab150077.html |
| Mounting medium with DAPI | abcam | ab104139 | GR3299210-7 | https://www.abcam.com/products/reagents/mounting-medium-with-dapi-aqueous-fluoroshield-ab104139.html |

Western blot

Western blot
| Materials | Vendor or Source | Catalog # | Lot # | Persistent ID / URL |
| --- | --- | --- | --- | --- |
| RIPA buffer | Cell Signaling Technology | 9803 | 35 | https://www.cellsignal.com/products/buffers-dyes/cell-lysis-buffer-10x/9803 |
| Protease inhibitor cocktail | MilliporeSigma | P8340-1ML | 126M4015 | https://www.sigmaaldrich.com/US/en/product/sigma/p8340 |
| Protein Assay Reagent A (DC assay) | Bio-Rad | 5000113 | 64343170 | https://www.bio-rad.com/en-us/SearchResults?search_api_fulltext=5000113 |
| Protein Assay Reagent B (DC assay) | Bio-Rad | 5000114 | 64327198 | https://www.bio-rad.com/en-us/SearchResults?search_api_fulltext=5000114 |
| Protein Assay Reagent S (DC assay) | Bio-Rad | 5000115 | 64327194 | https://www.bio-rad.com/en-us/SearchResults?search_api_fulltext=5000115 |
| Protein Standard II (DC assay) | Bio-Rad | 5000007 | 64368287 | https://www.bio-rad.com/en-us/SearchResults?search_api_fulltext=5000007 |
| Trans-Blot Turbo Mini 0.2 µm PVDF Transfer Packs | Bio-Rad | 1704156 | 64526914 | https://www.bio-rad.com/en-us/SearchResults?search_api_fulltext=1704156 |
| Trans-Blot Turbo Midi 0.2 µm PVDF Transfer Kit | Bio-Rad | 10026933 | BR20200810 | N/A |
| Goat Anti-Rabbit IgG Antibody (H+L), Peroxidase | Vector Laboratories | PI-1000-1 | ZG1009 | https://vectorlabs.com/products/antibodies/peroxidase-goat-anti-rabbit-igg |
| Goat Anti-Mouse IgG peroxidase antibody produced | MilliporeSigma | A2554 | 175122 | https://www.sigmaaldrich.com/US/en/product/sigma/a2554 |
| Protein Deglycosylation Mix II | New England Biolabs | P6044 | 10160304 | https://www.neb.com/protocols/2016/07/29/reaction-protocols-for-protein-deglycosylation-mix-ii-p6044 |
| Protein Deglycosylation Mix II (Mix Buffer 2) | New England Biolabs | P6044 | 10160306 | https://www.neb.com/protocols/2016/07/29/reaction-protocols-for-protein-deglycosylation-mix-ii-p6044 |

Cultured Cells

Cultured Cells
| Materials | Vendor or Source | Catalog # | Lot # | Persistent ID / URL |
| --- | --- | --- | --- | --- |
| HepG2 cell | ATCC | HB-8065 | N/A | https://www.atcc.org/products/hb-8065 |
| DMEM (high glucose) | Gibco | 12100046 | N/A | https://www.thermofisher.com/order/catalog/product/12100046?SID=srch-hj-12100046 |
| FBS | R&D | S12450H | L21260 | https://www.rndsystems.com/products/fetal-bovine-serum-optima-heat-inactivated_s12450h |
| Penicillin-Streptomycin | Gibco | 15070063 | 2441413 | https://www.thermofisher.com/order/catalog/product/15070063 |
| GlutaMax | Gibco | 35050061 | 2277245 | https://www.thermofisher.com/order/catalog/product/35050061 |
| 4-well chamber slide | Thermo Scientific | 154526 | 02022341 | https://www.thermofisher.com/order/catalog/product/154526 |

### Study designing following the ARRIVE Essential 10

Supplemental Figure 1

Supplemental Figure 1
| Groups | Sex | Age (week) | Number (prior to experiment) | Number (termination) | Littermates (Yes/No) |
| --- | --- | --- | --- | --- | --- |
| hepAGT<sup>+/+</sup> mice injected with AAV.null | M | 7-14 | 8 | 8 | Yes |
| hepAGT<sup>-/-</sup> mice injected with AAV.null | M | 7-14 | 6 | 6 |  |
| hepAGT<sup>-/-</sup> mice injected with AAV.mAGT | M | 7-14 | 8 | 8 |  |

Figure 1C-E and Supplemental Figures 2, 4, and 12

Figure 1C-E and Supplemental Figures 2, 4, and 12
| Groups | Sex | Age (week) | Number (prior to experiment) | Number (termination) | Littermates (Yes/No) |
| --- | --- | --- | --- | --- | --- |
| hepAGT<sup>-/-</sup> mice injected with AAV.null | M | 8-22 | 9 | 9 | Yes |
| hepAGT<sup>-/-</sup> mice injected with AAV.mAGT | M | 8-22 | 9 | 9 |  |
| hepAGT<sup>-/-</sup> mice injected with AAV.loop-Mut | M | 8-22 | 10 | 10 |  |

Figure 1G-I and Supplemental Figures 3, 5, and 12

Figure 1G-I and Supplemental Figures 3, 5, and 12
| Groups | Sex | Age (week) | Number (prior to experiment) | Number (termination) | Littermates (Yes/No) |
| --- | --- | --- | --- | --- | --- |
| hepAGT<sup>-/-</sup> mice injected with AAV.null | M | 6-20 | 6 | 6 | Yes |
| hepAGT<sup>-/-</sup> mice injected with AAV.mAGT | M | 6-20 | 6 | 6 |  |
| hepAGT<sup>-/-</sup> mice injected with AAV.βsheet-Mut | M | 6-20 | 6 | 6 |  |

Figure 2A-F, Supplemental Figures 6 – 8 and 10 – 12

Figure 2A-F, Supplemental Figures 6 – 8 and 10 - 12
| Groups | Sex | Age (week) | Number (prior to experiment) | Number (termination) | Littermates (Yes/No) |
| --- | --- | --- | --- | --- | --- |
| hepAGT-/- mice injected with AAV.null | M | 6-16 | 12 | 12 | Yes |
| hepAGT-/- mice injected with AAV.mAGT | M | 6-16 | 12 | 12 |  |
| hepAGT-/- mice injected with AAV.loop/βsheet-Mut | M | 6-16 | 13 | 13 |  |

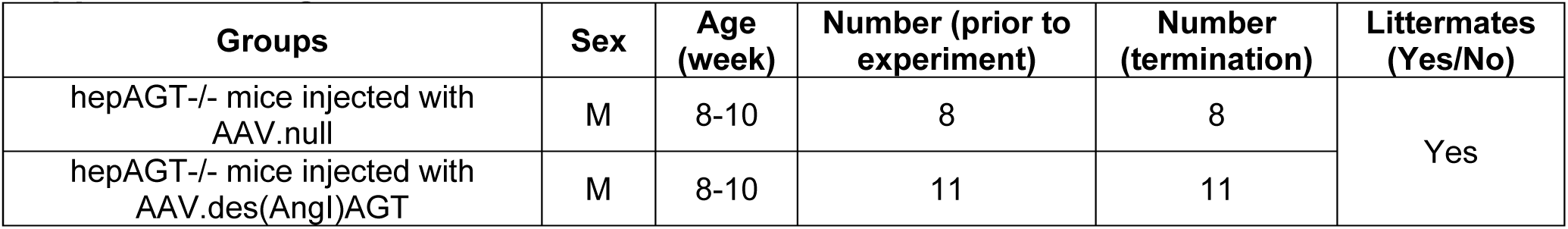
Supplemental Figure 9

ARRIVE Guidelines Checklists

ARRIVE Guidelines Checklists
| Item | Application |
| --- | --- |
| Ethics | Approved by the University of Kentucky IACUC (2018-2968). |
| Sex | This study included only male mice because of the modest effects of AAVs encoding AGT on plasma AGT concentrations in female mice.<sup>3</sup> |
| Inclusion criteria | Body weight > 18 g |
| Exclusion criteria | 1. Body weight ≤ 18 g<br>2. Medical cases reported by a veterinarian |
| Sample size | Described in each figure legend. |
| Sample size calculation | Based on the previous report,<sup>3</sup> studies were designed to include at least n=5 per group for evaluating the primary endpoints. |
| Endpoint | 1. Plasma AGT concentrations.<br>2. Systolic blood pressure.<br>3. Percentage lesion area of atherosclerosis in the aortic arch. |
| Randomization | Study mice were numbered randomly and allocated to study groups based on the numbers. |
| Blinding | All experimental data were verified by an independent investigator blinded to the study group information. |
| Statistical analysis | SigmaPlot version 14.5 or 15.0 (SYSTAT Software Inc., CA). |
| Statistical method | Described in each figure legend |
| Data availability | All numerical data are available in Supplemental Excel File I. All raw data and analytical methods are available from the corresponding authors upon appropriate request. |

---

**Supplemental Figure 1.**
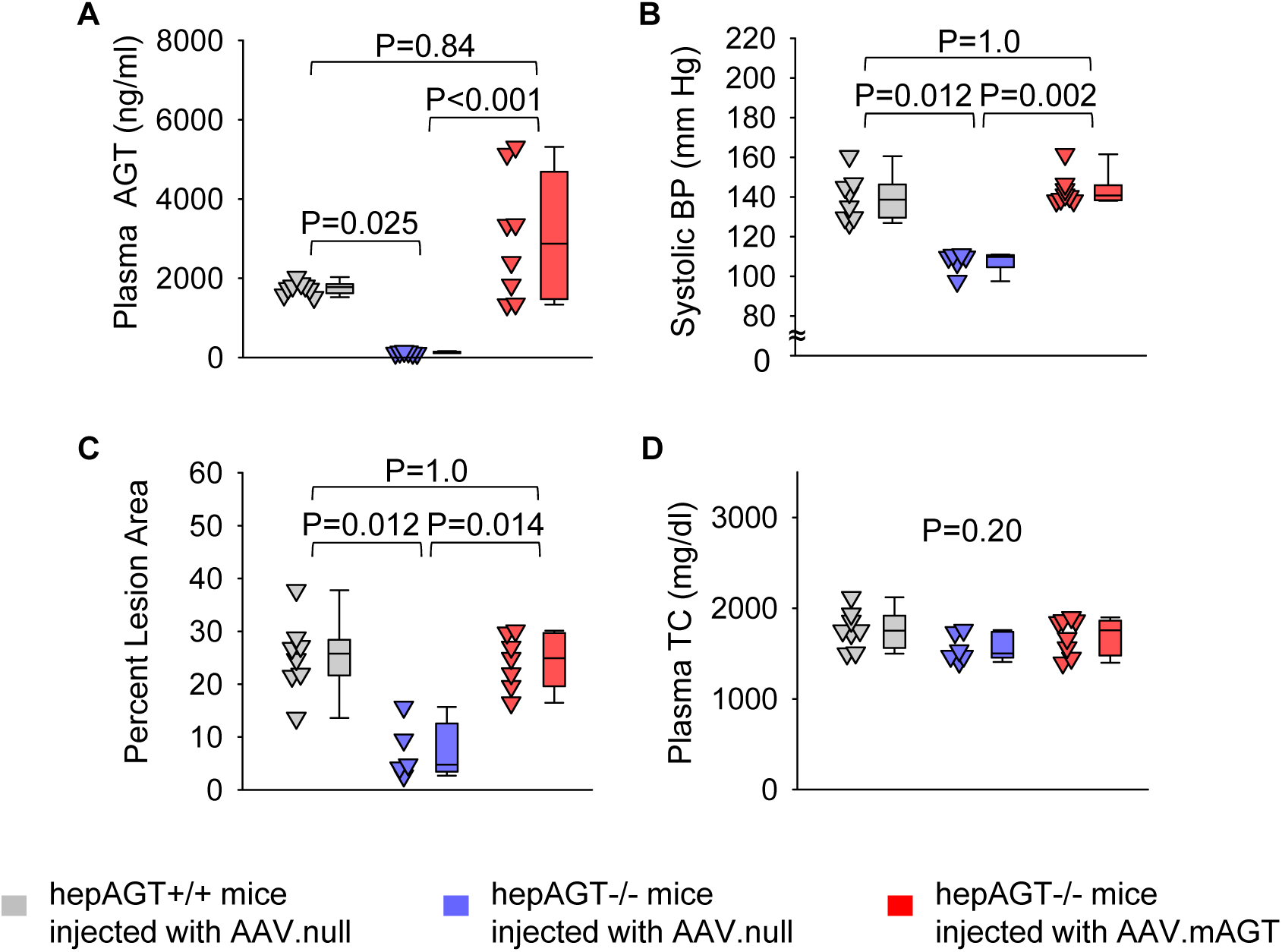
Population of wild-type AGT increased plasma AGT concentrations in hepAGT-/– mice to the comparable magnitude of hepAGT+/+ mice. All study mice were male in an LDL receptor –/– background and fed Western diet for 12 weeks. **(A)** Plasma AGT concentrations, **(B)** Systolic blood pressure (BP), **(C)** percent atherosclerotic lesion areas in the ascending aorta and the aortic arch regions, and **(D)** plasma total cholesterol concentrations of hepAGT+/+ mice administered AAV.null or hepAGT-/– mice administered AAV.null or AAV.mAGT (n = 5-8/group). P values were determined by Kruskal-Wallis one-way ANOVA on Ranks followed by Dunn’s test. AAV.null: AAV containing a null vector; AAV.mAGT: AAV encoding mouse wild-type AGT.

**Supplemental Figure 2.**
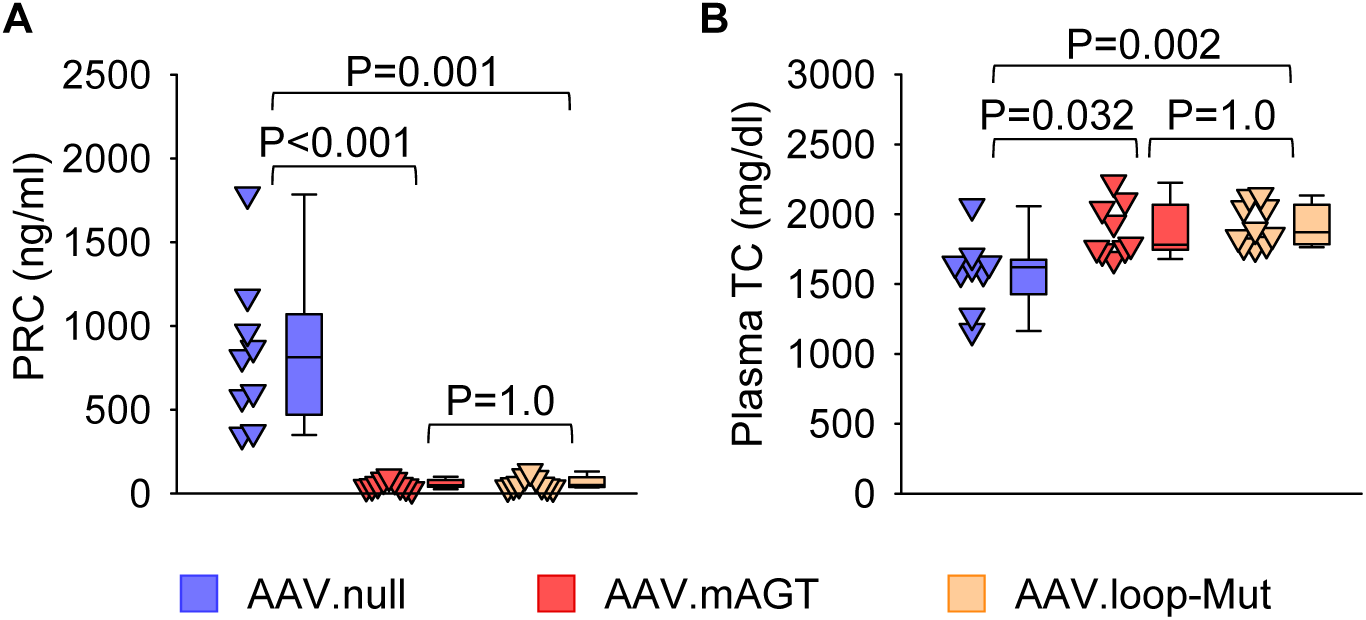
Mutations of the conserved sequences in the loop region of AGT had no effects on plasma renin concentrations (PRC) and total cholesterol (TC) concentrations. All study mice were male in an LDL receptor –/– background and fed Western diet for 12 weeks. **(A)** PRC and **(B)** plasma TC concentrations of hepAGT-/– mice administered AAV.null, AAV.mAGT, or AAV.loop-Mut (n = 9-10/group). P values were determined by Kruskal-Wallis one-way ANOVA on Ranks followed by Dunn’s test. AAV.null: AAV containing a null vector; AAV.mAGT: AAV encoding mouse wild-type AGT; AAV.loop-Mut: AAV encoding mouse AGT with mutations of conserved sequences in the loop region.

**Supplemental Figure 3.**
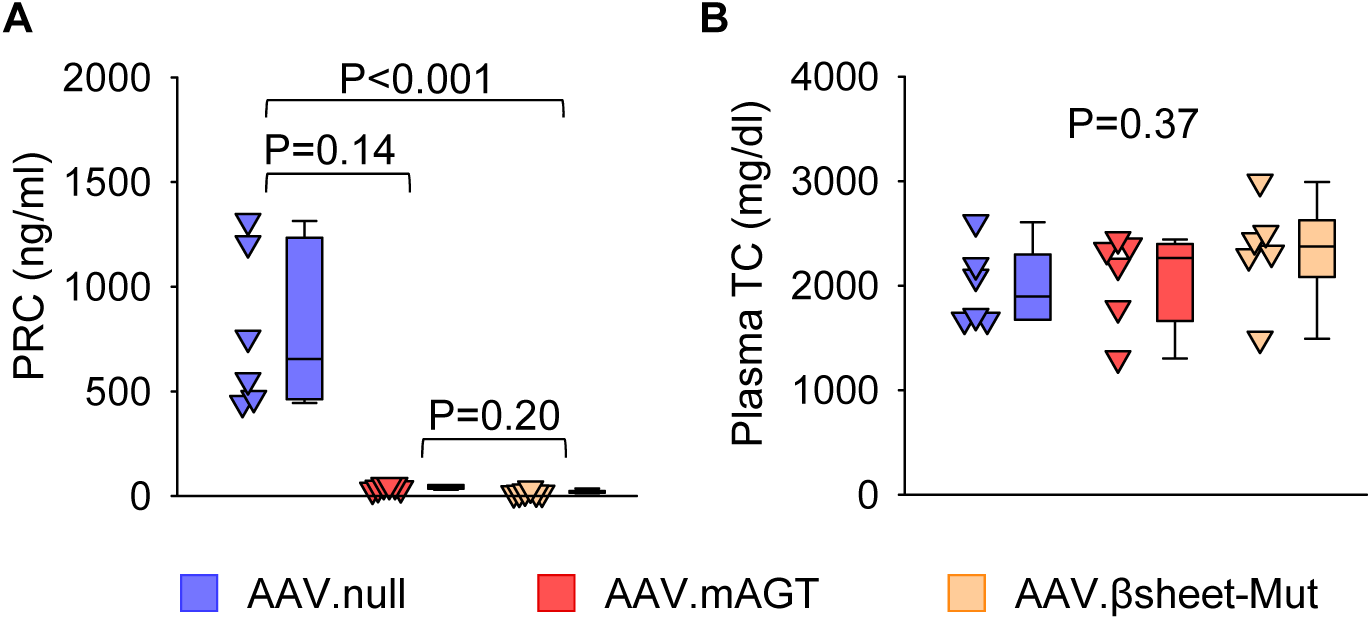
Mutations of the conserved sequences in the β-sheet region of AGT had no effects on plasma renin concentrations (PRC) and total cholesterol (TC) concentrations. All study mice were male in an LDL receptor –/– background and fed Western diet for 12 weeks. **(A)** PRC and **(B)** plasma TC concentrations of hepAGT–/– mice administered AAV.null, AAV.mAGT, or AAV.βsheet-Mut (n = 6/group). P values were determined by Kruskal-Wallis one-way ANOVA on Ranks followed by Dunn’s test. AAV.null: AAV containing a null vector; AAV.mAGT: AAV encoding mouse wild-type AGT; AAV.βsheet-Mut: AAV encoding mouse AGT with mutations of conserved sequences in the β-sheet region.

**Supplemental Figure 4.**
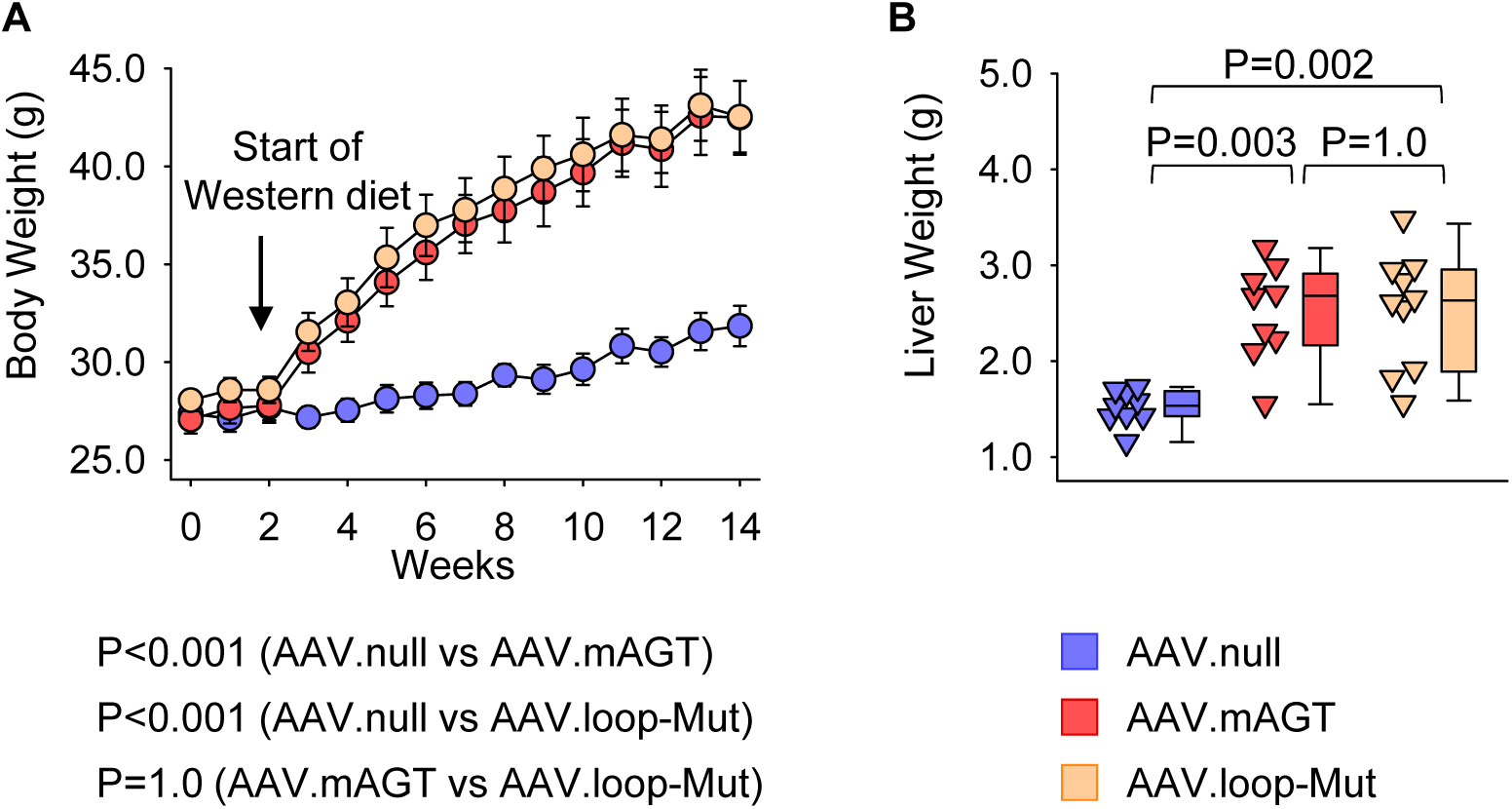
Population of AGT with mutations of conserved sequences in the loop led to diet-induced body weight and liver weight gain in hepAGT-/– mice. All study mice were male in an LDL receptor –/– background and fed Western diet for 12 weeks. **(A)** Body weight gain during the study, and **(B)** liver weight at termination of hepAGT-/– mice administered AAV.null, AAV.mAGT, or AAV.loop-Mut (n = 9-10/group). P values were determined using a linear mixed-effects model followed by Bonferroni test (A) or Kruskal-Wallis one-way ANOVA on Ranks followed by Dunn’s test (B). AAV.null: AAV containing a null vector; AAV.mAGT: AAV encoding mouse wild-type AGT; AAV.loop-Mut: AAV encoding mouse AGT with mutations of conserved sequences in the loop region.

**Supplemental Figure 5.**
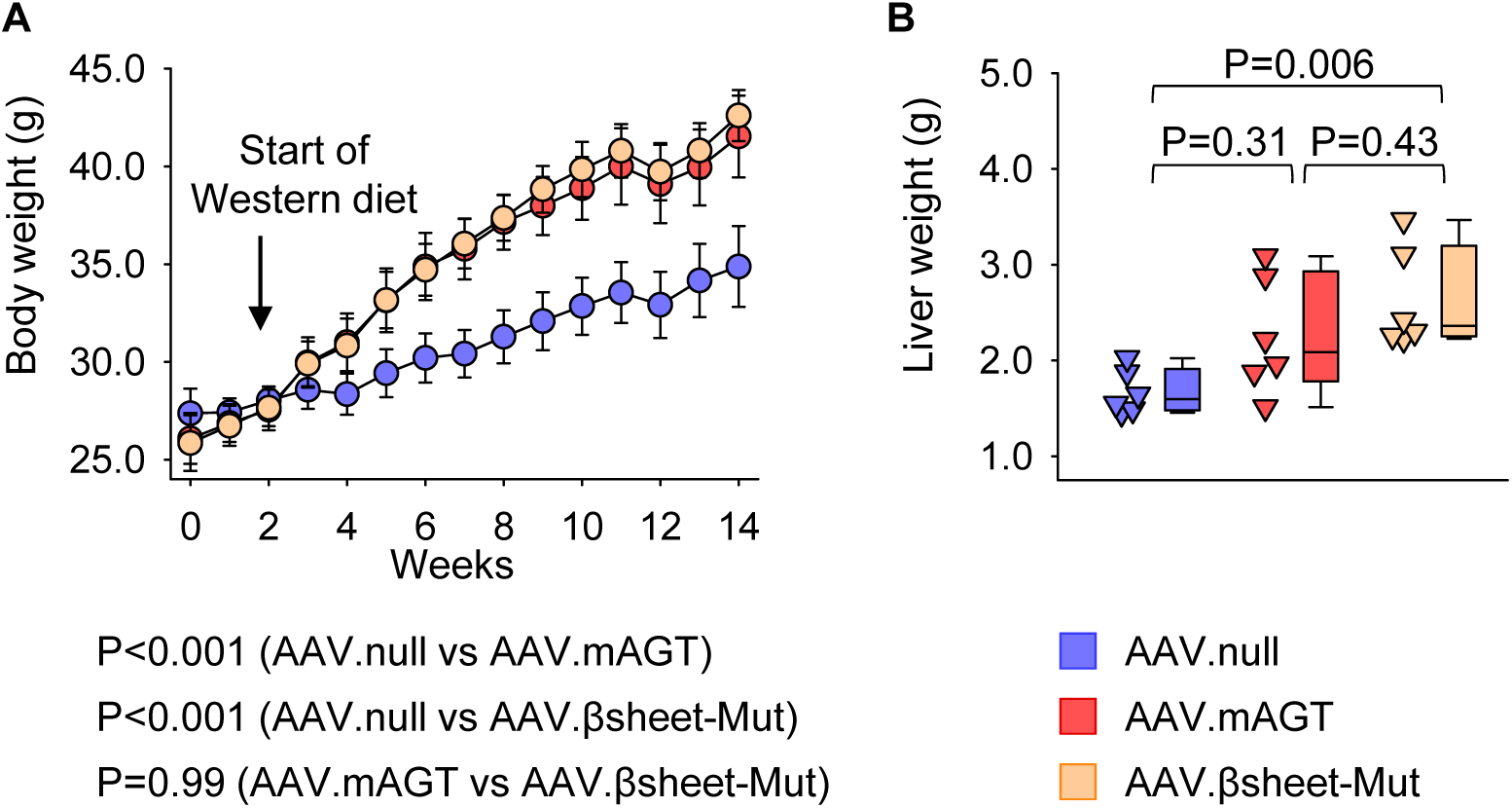
Population of AGT with mutations of conserved sequences in the β-sheet region led to diet-induced body weight and liver weight gain in hepAGT-/– mice. All study mice were male in an LDL receptor –/– background and fed Western diet for 12 weeks. **(A)** Body weight gain during the study, and **(B)** liver weight at termination of hepAGT-/– mice administered AAV.null, AAV.mAGT, or AAV.βsheet-Mut (n = 6/group). P values were determined using a linear mixed-effects model followed by Bonferroni test (A) or Kruskal-Wallis one-way ANOVA on Ranks followed by Dunn’s test (B). AAV.null: AAV containing a null vector; AAV.mAGT: AAV encoding mouse wild-type AGT; AAV.βsheet-Mut: AAV encoding mouse AGT with mutations of conserved sequences in the β-sheet region.

**Supplemental Figure 6.**
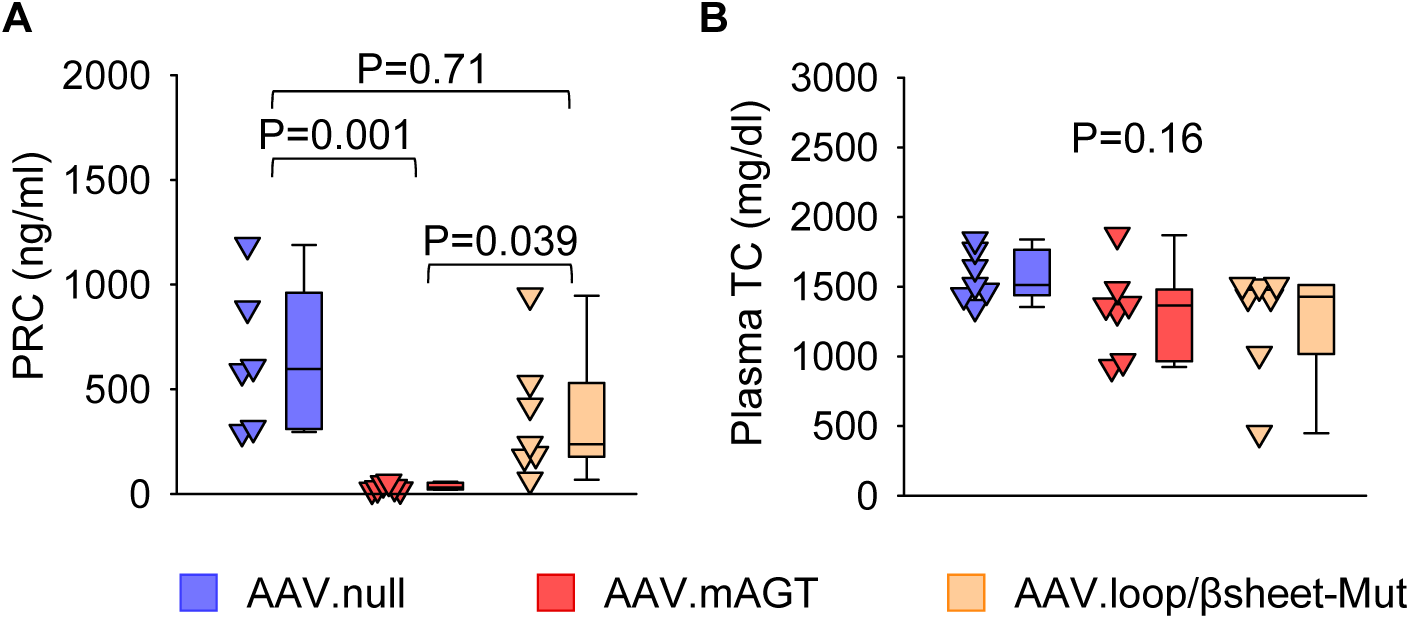
Mutations of the conserved sequences in both the loop and β– sheet regions of AGT did not change plasma renin and total cholesterol concentrations in hepAGT-/– mice. All study mice were male in an LDL receptor –/– background and fed Western diet for 12 weeks. **(A)** PRC and **(B)** plasma TC concentrations of hepAGT-/– mice administered AAV.null, AAV.mAGT, or AAV.loop/βsheet-Mut (n= 6-7/group). P values were determined by Kruskal-Wallis one-way ANOVA on Ranks followed by Dunn’s test. AAV.null: AAV containing a null vector; AAV.mAGT: AAV encoding mouse wild-type AGT; AAV.loop/βsheet-Mut: AAV encoding mouse AGT with mutations of conserved sequences in both the loop and β-sheet regions; PRC: plasma renin concentrations; TC: total cholesterol.

**Supplemental Figure 7.**
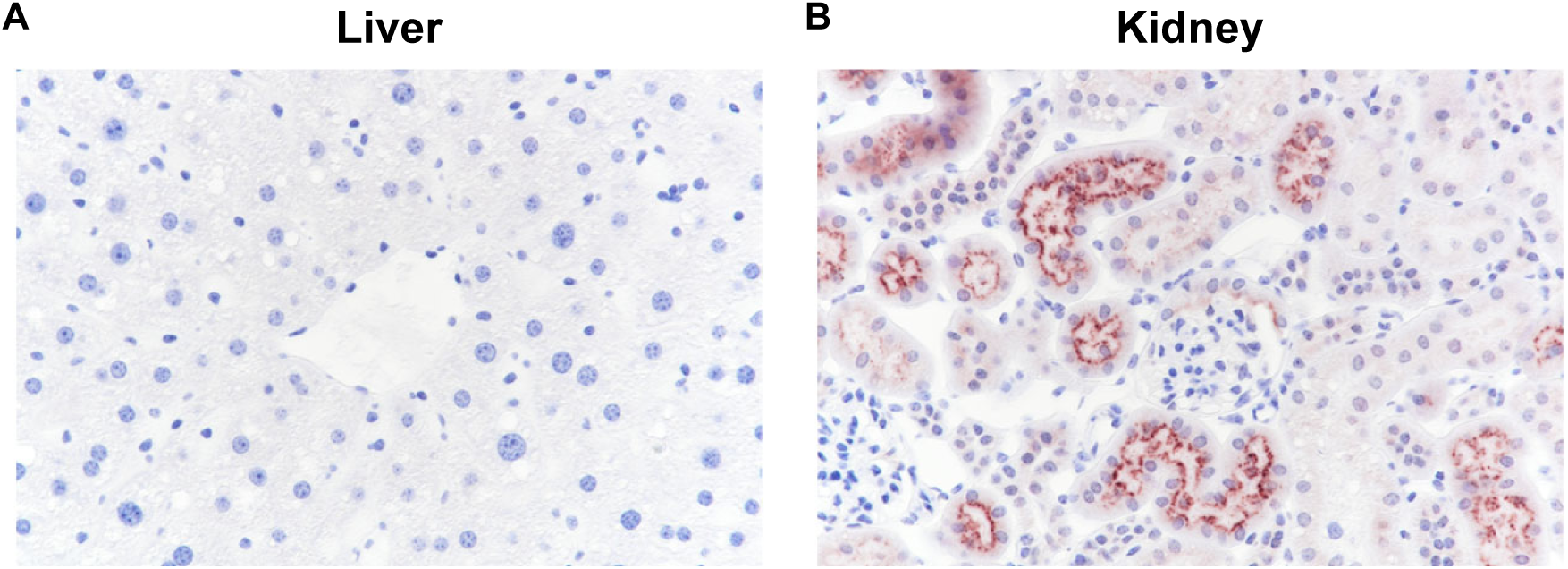
Immunostaining of AGT in liver and kidney from hepAGT+/+ mice. Tissues were obtained from male mice in an LDL receptor –/– background and fed Western diet for 12 weeks. Representative images of immunostaining for AGT in **(A)** liver and **(B)** kidney sections.

**Supplemental Figure 8.**
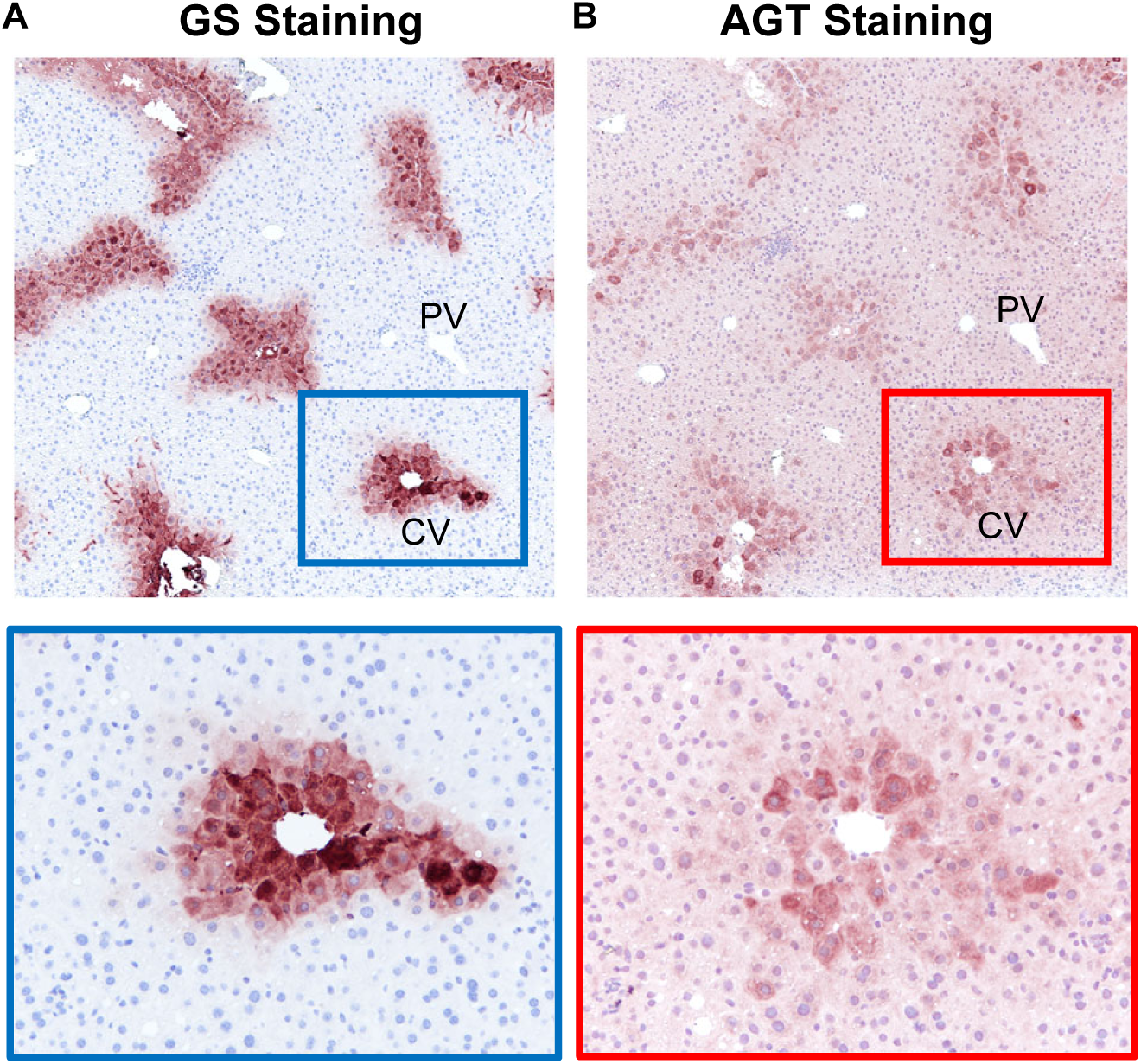
Immunostaining of glutamine synthetase (GS) and AGT in mouse liver sections. Immunostaining of **(A)** glutamine synthetase and **(B)** AGT in liver sections obtained from hepAGT-/– male mice administered AAV.loop/βsheet-Mut. High-magnification 40× images were represented in blue and red boxes of the bottom panel. AAV.loop/βsheet-Mut: AAV encoding mouse AGT with mutations of conserved sequences in both the loop and β-sheet regions; CV: central vein; PV: portal vein.

**Supplemental Figure 9.**
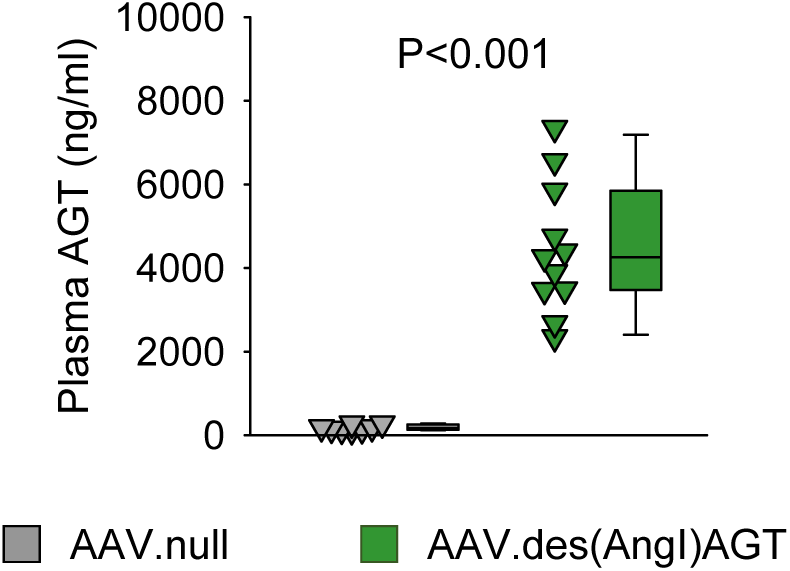
Population of des(AngI)AGT increased plasma AGT concentrations in hepAGT-/– mice. All study mice were male male in an LDL receptor –/– background and fed Western diet for 12 weeks. Plasma AGT concentrations were measured 14 weeks after administration of AAV.des(AngI)AGT (n = 8-11/group). P value was determined by Mann-Whitney Rank Sum Test. AAV.null: AAV containing a null vector; AAV.des(AngI)AGT: AAV encoding mouse des(AngI)AGT.

**Supplemental Figure 10.**
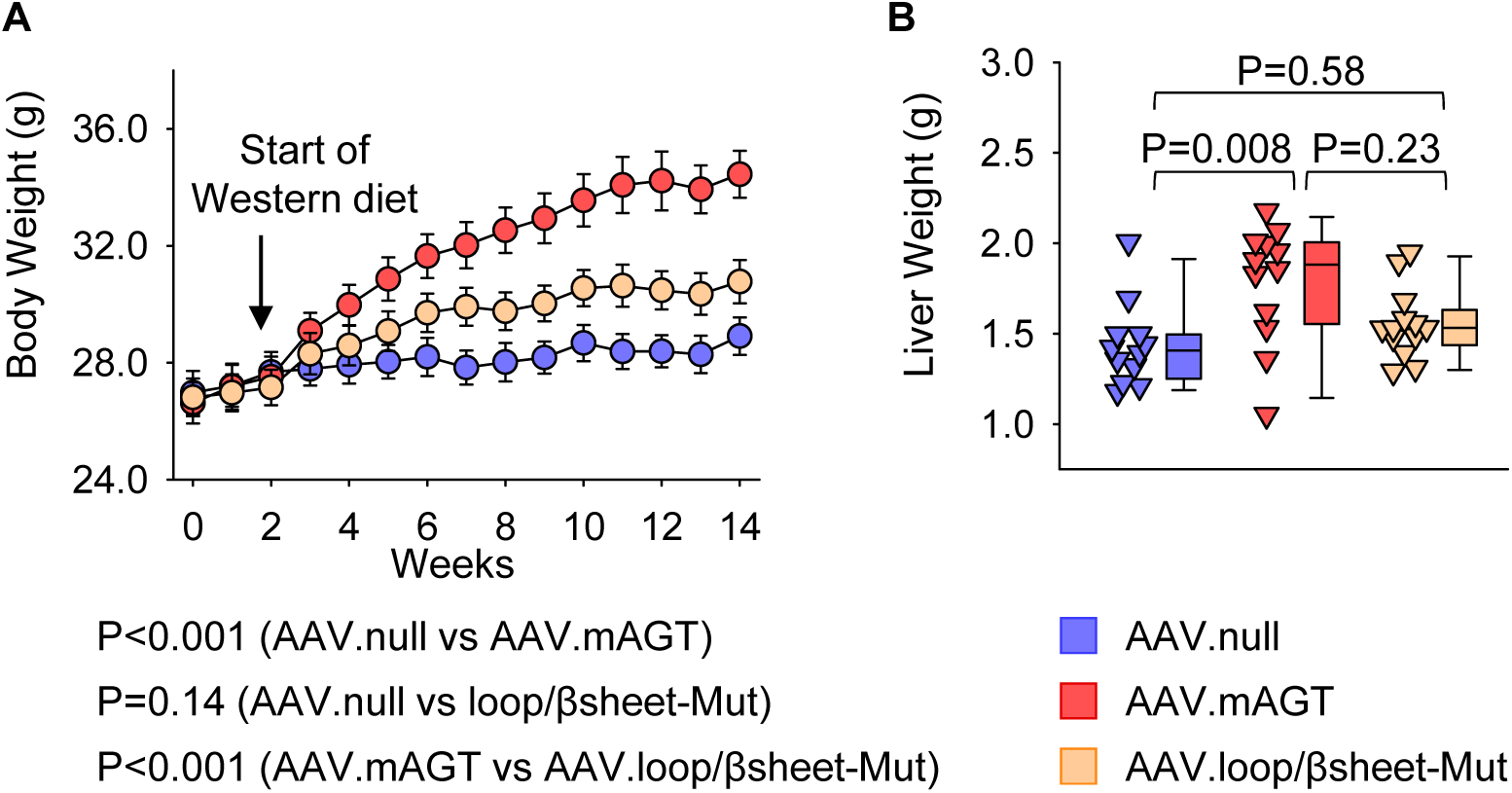
Population of AGT with mutations of conserved sequences in both the loop and β-sheet did not affect diet-induced body weight and liver weight gain in hepAGT-/– mice. All study mice were male in an LDL receptor –/– background and fed Western diet for 12 weeks. **(A)** Body weight gain during the study, and **(B)** liver weight at termination of hepAGT-/– mice administered AAV.null, AAV.mAGT, or AAV.loop/βsheet-Mut (n = 12-13/group). P values were determined using a linear mixed-effects model followed by Bonferroni test (A) or Kruskal-Wallis one-way ANOVA on Ranks followed by Dunn’s test (B). AAV.null: AAV containing a null vector; AAV.mAGT: AAV encoding mouse wild-type AGT; AAV.loop/βsheet-Mut: AAV encoding mouse AGT with mutations of conserved sequences in both the loop and β-sheet regions.

**Supplemental Figure 11.**
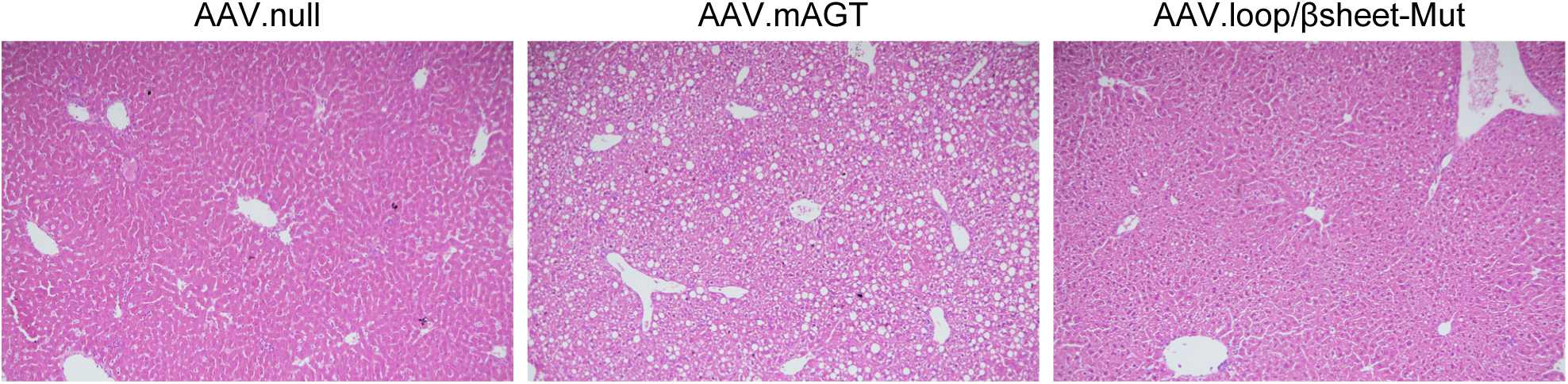
Histological assessment of liver steatosis. Representative images of hematoxylin and eosin staining of liver samples from mice administered AAV.null, AAV.mAGT, or AAV.loop/βsheet-Mut. AAV.null: AAV containing a null vector; AAV.mAGT: AAV encoding mouse wild-type AGT; AAV.loop/βsheet-Mut: AAV encoding mouse AGT with mutations of conserved sequences in both the loop and β-sheet regions.

**Supplemental Figure 12.**
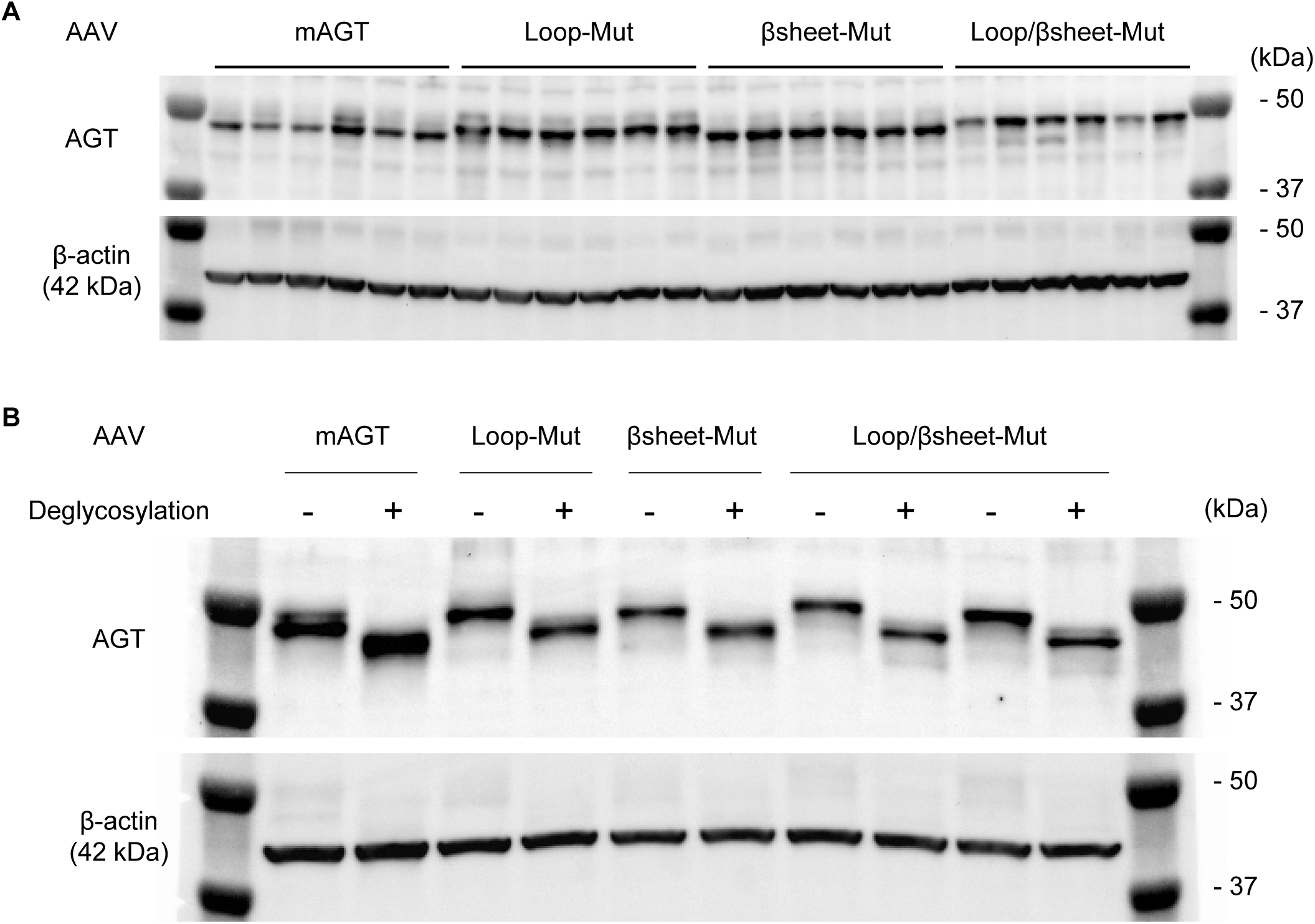
Mutations in either or both the loop and the β-sheet regions of AGT in liver did not impact the glycosylation of AGT. (**A**) Western blot for AGT using proteins of liver tissues from hepAGT-/– mice administered AAV.mAGT, AAV.loop-Mut, AAV.βsheet-Mut, or AAV.loop/βsheet-Mut (n = 6/group). **(B)** Western blot for AGT using proteins of liver samples with or without deglycosylation. mAGT: mouse wild-type AGT; loop-Mut: mouse AGT with mutations of conserved sequences in the loop region; βsheet-Mut: mouse AGT with mutations of conserved sequences in the β-sheet region; loop/βsheet-Mut: mouse AGT with mutations of conserved sequences in both the loop and β-sheet regions.

## REFERENCES

1. Daugherty A, Manning MW and Cassis LA. Angiotensin II promotes atherosclerotic lesions and aneurysms in apolipoprotein E-deficient mice. J Clin Invest. 2000;105:1605–1612.

2. Ruiz-Ortega M, Lorenzo O, Rupérez M, Esteban V, Suzuki Y, Mezzano S, Plaza JJ and Egido J. Role of the renin-angiotensin system in vascular diseases: expanding the field. Hypertension. 2001;38:1382–1387.

3. Weiss D, Sorescu D and Taylor WR. Angiotensin II and atherosclerosis. Am J Cardiol. 2001;87:25C–32C.

4. Weiss D, Kools JJ and Taylor WR. Angiotensin II-induced hypertension accelerates the development of atherosclerosis in apoE-deficient mice. Circulation. 2001;103:448–454.

5. Rajagopalan S, Kurz S, Münzel T, Tarpey M, Freeman BA, Griendling KK and Harrison DG. Angiotensin II-mediated hypertension in the rat increases vascular superoxide production via membrane NADH/NADPH oxidase activation. Contribution to alterations of vasomotor tone. J Clin Invest. 1996;97:1916–1923.

6. Mehta PK and Griendling KK. Angiotensin II cell signaling: physiological and pathological effects in the cardiovascular system. Am J Physiol Cell Physiol. 2007;292:C82–97.

7. Clouston WM, Evans BA, Haralambidis J and Richards RI. Molecular cloning of the mouse angiotensinogen gene. Genomics. 1988;2:240–248.

8. Lynch KR and Peach MJ. Molecular biology of angiotensinogen. Hypertension. 1991;17:263–269.

9. Wu C, Lu H, Cassis LA and Daugherty A. Molecular and pathophysiological features of angiotensinogen: a mini review. N Am J Med Sci (Boston). 2011;4:183–190.

10. Lu H, Cassis LA, Kooi CW and Daugherty A. Structure and functions of angiotensinogen. Hypertens Res. 2016;39:492–500.

11. Célérier J, Cruz A, Lamandé N, Gasc JM and Corvol P. Angiotensinogen and its cleaved derivatives inhibit angiogenesis. Hypertension. 2002;39:224–228.

12. Corvol P, Lamandé N, Cruz A, Celerier J and Gasc JM. Inhibition of angiogenesis: a new function for angiotensinogen and des(angiotensin I)angiotensinogen. Curr Hypertens Rep. 2003;5:149–154.

13. Lu H, Wu C, Howatt DA, Balakrishnan A, Moorleghen JJ, Chen X, Zhao M, Graham MJ, Mullick AE, Crooke RM, Feldman DL, Cassis LA, Vander Kooi CW and Daugherty A. Angiotensinogen exerts effects independent of Angiotensin II. Arterioscler Thromb Vasc Biol. 2016;36:256–265.

14. Tao XR, Rong JB, Lu HS, Daugherty A, Shi P, Ke CL, Zhang ZC, Xu YC and Wang JA. Angiotensinogen in hepatocytes contributes to Western diet-induced liver steatosis. J Lipid Res. 2019;60:1983–1995.

15. Zhou A, Carrell RW, Murphy MP, Wei Z, Yan Y, Stanley PL, Stein PE, Broughton Pipkin F and Read RJ. A redox switch in angiotensinogen modulates angiotensin release. Nature. 2010;468:108–111.

16. Wu C, Xu Y, Lu H, Howatt DA, Balakrishnan A, Moorleghen JJ, Vander Kooi CW, Cassis LA, Wang JA and Daugherty A. Cys18-Cys137 disulfide bond in mouse angiotensinogen does not affect AngII-dependent functions in vivo. Hypertension. 2015;65:800–805.

17. Ye F, Wang Y, Wu C, Howatt DA, Wu CH, Balakrishnan A, Mullick AE, Graham MJ, Danser AHJ, Wang J, Daugherty A and Lu HS. Angiotensinogen and megalin interactions contribute to atherosclerosis-Brief report. Arterioscler Thromb Vasc Biol. 2019;39:150–155.

18. Matsusaka T, Niimura F, Shimizu A, Pastan I, Saito A, Kobori H, Nishiyama A and Ichikawa I. Liver angiotensinogen is the primary source of renal angiotensin II. J Am Soc Nephrol. 2012;23:1181–1189.

19. Yiannikouris F, Wang Y, Shoemaker R, Larian N, Thompson J, English VL, Charnigo R, Su W, Gong M and Cassis LA. Deficiency of angiotensinogen in hepatocytes markedly decreases blood pressure in lean and obese male Mice. Hypertension. 2015;66:836–842.

20. Wu CH, Wu C, Howatt DA, Moorleghen JJ, Cassis LA, Daugherty A and Lu HS. Two amino acids proximate to the renin cleavage site of human angiotensinogen do not affect blood pressure and atherosclerosis in mice. Arterioscler Thromb Vasc Biol. 2020;40:2108–2113.

21. Ye D, Wu C, Chen H, Liang CL, Howatt DA, Franklin MK, Moorleghen JJ, Tyagi SC, Uijl E, Danser AHJ, Sawada H, Daugherty A and Lu HS. Fludrocortisone induces aortic pathologies in mice. Biomolecules. 2022;12:825.

22. Bell P, Wang L, Gao G, Haskins ME, Tarantal AF, McCarter RJ, Zhu Y, Yu H and Wilson JM. Inverse zonation of hepatocyte transduction with AAV vectors between mice and non-human primates. Mol Genet Metab. 2011;104:395–403.

23. Daugherty A, Rateri D, Hong L and Balakrishnan A. Measuring blood pressure in mice using volume pressure recording, a tail-cuff method. J Vis Exp. 2009.

24. Daugherty A, Tall AR, Daemen M, Falk E, Fisher EA, García-Cardeña G, Lusis AJ, Owens AP, 3rd, Rosenfeld ME and Virmani R. Recommendation on design, execution, and reporting of animal atherosclerosis studies: A scientific statement from the American Heart Association. Arterioscler Thromb Vasc Biol. 2017;37:e131–e157.

25. Chen H, Howatt DA, Franklin MK, Amioka N, Sawada H, Daugherty A and Lu HS. A mini-review on quantification of atherosclerosis in hypercholesterolemic mice. Glob Transl Med. 2022;1:72.

26. Viotti C. ER to Golgi-dependent protein secretion: the conventional pathway. Methods Mol Biol. 2016;1459:3–29.

27. Benham AM. Protein secretion and the endoplasmic reticulum. Cold Spring Harb Perspect Biol. 2012;4:a012872.

28. Gomez-Navarro N and Miller E. Protein sorting at the ER-Golgi interface. J Cell Biol. 2016;215:769–778.

29. Ramazi S and Zahiri J. Posttranslational modifications in proteins: resources, tools and prediction methods. Database (Oxford). 2021;2021:baab012.

30. Gimenez-Roqueplo AP, Célérier J, Lucarelli G, Corvol P and Jeunemaitre X. Role of N-glycosylation in human angiotensinogen. J Biol Chem. 1998;273:21232–21238.

31. Chen SJ, Sanmiguel J, Lock M, McMenamin D, Draper C, Limberis MP, Kassim SH, Somanathan S, Bell P, Johnston JC, Rader DJ and Wilson JM. Biodistribution of AAV8 vectors expressing human low-density lipoprotein receptor in a mouse model of homozygous familial hypercholesterolemia. Hum Gene Ther Clin Dev. 2013;24:154–160.

32. Yan Y, Zhou A, Carrell RW and Read RJ. Structural basis for the specificity of renin-mediated angiotensinogen cleavage. J Biol Chem. 2019;294:2353–2364.

33. Lomas DA and Mahadeva R. Alpha1-antitrypsin polymerization and the serpinopathies: pathobiology and prospects for therapy. J Clin Invest. 2002;110:1585–1590.

34. Wu CH, Mohammadmoradi S, Chen JZ, Sawada H, Daugherty A and Lu HS. Renin-angiotensin system and cardiovascular functions. Arterioscler Thromb Vasc Biol. 2018;38:e108–e116.

35. Morgan ES, Tami Y, Hu K, Brambatti M, Mullick AE, Geary RS, Bakris GL and Tsimikas S. Antisense inhibition of angiotensinogen with IONIS-AGT-L(Rx): results of Phase 1 and Phase 2 studies. JACC Basic Transl Sci. 2021;6:485–496.

36. Uijl E, Ye D, Ren L, Mirabito Colafella KM, van Veghel R, Garrelds IM, Lu HS, Daugherty A, Hoorn EJ, Nioi P, Foster D and Danser AHJ. Conventional vasopressor and vasopressor-sparing strategies to counteract the blood pressure-lowering effect of small interfering RNA targeting angiotensinogen. J Am Heart Assoc. 2022;11:e026426.

## REFERENCES

1. Zhou A, Carrell RW, Murphy MP, Wei Z, Yan Y, Stanley PL, Stein PE, Broughton Pipkin F and Read RJ. A redox switch in angiotensinogen modulates angiotensin release. Nature. 2010;468:108–111.

2. Cotugno G, Annunziata P, Barone MV, Karali M, Banfi S and Auricchio A. Impact of age at administration, lysosomal storage, and transgene regulatory elements on AAV2/8-mediated rat liver transduction. PLoS One. 2012;7:e33286.

3. Wu CH, Wu C, Howatt DA, Moorleghen JJ, Cassis LA, Daugherty A and Lu HS. Two amino acids proximate to the renin cleavage site of human angiotensinogen do not affect blood pressure and atherosclerosis in mice. Arterioscler Thromb Vasc Biol. 2020;40:2108–2113.

